# The BAF chromatin remodeling subunit ARID1B coordinates cell-cycle exit and motor neuron differentiation

**DOI:** 10.64898/2026.10.24.761583

**Authors:** Alyssa Ialongo Del Prete, Ssu-Yu Yeh, Kaitlin N. Montanera, Ravneet Jaura, Ho Sung Rhee

**Author notes:** Corresponding author: Ho Sung Rhee.

## Abstract

Neuronal differentiation requires precise regulation of transcriptional and epigenetic networks by BAF (BRG1/BRM-associated factor) chromatin remodeling complexes. Mutations in *Arid1b*, encoding a core BAF subunit, are associated with motor dysfunction, yet the role of ARID1B in motor neuron specification remains unclear. Using *Arid1b* knockout mouse embryonic stem cells, we show that ARID1B is not required for neural progenitor specification but is essential for the transition to postmitotic motor neurons. *Arid1b* knockout impairs cell-cycle exit and neurite outgrowth, reducing neuronal gene expression while maintaining early developmental and cell-cycle gene expression. Mechanistically, ARID1B-containing BAF complexes are recruited to neuronal gene promoters enriched for active chromatin signatures and the neuron-associated transcription factor NRF1 binding. The histone acetyltransferase p300 preferentially occupies ARID1B/BRG1 co-bound regions enriched with histone H3 lysine 27 acetylation (H3K27ac), and loss of ARID1B alters H3K27ac enrichment associated with neuronal gene expression. Together, these findings establish ARID1B as a critical regulator coordinating the developmental switch from cell-cycle progression to motor neuron differentiation.

## Introduction

Neuronal differentiation requires precise spatiotemporal regulation of cell type-specific gene expression programs by neuronal transcription factors (TFs), which drive neural lineage commitment, neuronal maturation, and function^1,2^. Thus, the ability of neuronal TFs to access the *cis*-regulatory regions of their target genes, including promoters and enhancers, is crucial for gene regulation. The mammalian genome is organized into regions of densely packed, transcriptionally inactive heterochromatin and loosely packed, transcriptionally active euchromatin^3^. Since most TFs are unable to access DNA within compacted chromatin, chromatin remodeling dynamically alters chromatin structure to ensure that *cis*-regulatory regions become accessible for TF binding at specific developmental stages^4^. ATP-dependent chromatin remodeling complexes facilitate this process through nucleosome sliding, eviction, and exchange, thereby regulating local chromatin structure^4^. Chromatin accessibility can also be modulated by covalent histone modifications, such as histone acetylation, which is generally associated with increased accessibility of *cis*-regulatory regions to TFs^5–7^.

The mammalian SWI/SNF complex, also referred to as the BAF (BRG1/BRM-associated factor) complex, is an ATP-dependent chromatin remodeling complex that regulates chromatin structure and gene expression during mammalian development^8,9^. BAF complexes are composed of 10-15 subunits, including the catalytic ATPase BRG1, and other subunits that can modulate genomic targeting in a cell type-specific manner^8–10^. The temporal reorganization of BAF subunit composition into neural progenitor BAF (npBAF) and neuronal BAF (nBAF) complexes is required for neuronal differentiation, as these complexes regulate gene expression programs associated with neural progenitors and differentiated neurons, respectively^11–13^. Mutations in the genes encoding npBAF and nBAF subunits are associated with a variety of central nervous system disorders in humans, highlighting the critical roles of these complexes in neurodevelopment^8^. However, the genome-wide functions of individual npBAF and nBAF subunits during neuronal differentiation remain incompletely understood.

The AT-rich interaction domain (ARID)-containing protein 1B (ARID1B) is a large, core subunit of both npBAF and nBAF complexes^14,15^. Pathogenic variants in *Arid1b* are estimated to account for approximately 1% of intellectual disability cases, of which approximately 50% are associated with autism spectrum disorder^16^. Additionally, *Arid1b* haploinsufficiency in humans and mice is associated with motor dysregulation^17,18^. Several studies have investigated the neurodevelopmental consequences of *Arid1b* deficiency by examining its function during cortical neuron differentiation^19–21^. Despite its importance in neurodevelopment, the mechanisms by which ARID1B regulates cell type-specific gene expression during neuronal differentiation remain incompletely understood.

One potential mechanism by which ARID1B may regulate neuronal gene expression is through the control of chromatin accessibility at cell type-specific *cis*-regulatory regions. BAF chromatin remodeling complexes rearrange nucleosomes to facilitate the activity of DNA-binding proteins at regulatory elements controlling genes required for neuronal cell identity and function^11,12^. Developmentally regulated *cis*-regulatory regions are dynamically remodeled during differentiation, and their activation is frequently associated with increased chromatin accessibility and enrichment of histone modifications such as histone H3 lysine 27 acetylation (H3K27ac). Whether and how ARID1B is linked to cell type-specific regulatory regions involved in the activation of neuronal genes through changes in chromatin accessibility and histone modifications remains unknown.

Given the motor abnormalities associated with *Arid1b* haploinsufficiency, motor neurons represent a particularly relevant neuronal population in which to investigate ARID1B function. Proper motor system function relies on the specification and maintenance of distinct motor neuron types^22^. Spinal motor neurons reside in the ventral horn of the spinal cord and transmit signals from the central nervous system to peripheral effector muscles^22^. These neurons extend long axonal projections that innervate target muscles and control movement. Their specification depends on a precisely coordinated transcriptional program directed by the proneural TF, Neurogenin-2 (NGN2), and the LIM homeobox TFs Islet1 (ISL1) and LHX3^23,24^. Disruption of these transcriptional programs may impair motor neuron specification and maintenance, potentially contributing to the motor dysfunction associated with *Arid1b* haploinsufficiency^17,18^. However, whether and how *Arid1b* mutations affect motor neuron specification remains poorly understood.

To address these questions, we generated an *Arid1b* knockout (KO) mouse embryonic stem (ES) cell line and used an *in vitro* differentiation system that efficiently directs mouse ES cells toward a spinal motor neuron fate^1^. We found that ARID1B is required for the activation of postmitotic neuronal genes and the repression of early developmental and cell-cycle genes during motor neuron differentiation. Furthermore, BAF complexes that contain ARID1B and BRG1 occupy a distinct subset of genomic regions characterized by active chromatin signatures. ARID1B-containing BAF complexes occupy *cis*-regulatory regions of target genes, where ARID1B loss is associated with altered H3K27ac and target gene expression. We further identified preferential p300 occupancy at H3K27ac-enriched regions co-bound by ARID1B and BRG1. Taken together, our findings identify ARID1B as a critical regulator of the transition from proliferating neural progenitors to postmitotic motor neurons and provide insight into how specialized BAF complexes coordinate transcriptional and epigenetic programs during motor neuron differentiation.

## Results

### Generation of an Arid1b knockout mouse ES cell line

To investigate the role of *Arid1b* mutation on motor neuron differentiation, we generated an *Arid1b* KO mouse ES cell line using CRISPR/Cas9-mediated genome editing. We introduced a homozygous frameshift deletion in exon 2 of *Arid1b*, resulting in a premature stop codon (**Fig. 1A**). This premature stop codon occurs upstream of the ARID domain and domain of unknown function 3518 (DUF3518), which mediate ARID1B-DNA and ARID1B-BRG1 interactions, respectively^25^. We confirmed that day 0 wild-type (WT) and *Arid1b* KO ES cell colonies exhibited similar alkaline phosphatase staining, indicative of ES cell pluripotency (**Fig. 1B**). We then differentiated day 0 WT and *Arid1b* KO ES cells into neural progenitors (day 4), nascent motor neurons (day 5), and maturing motor neurons (day 6) (**Fig. 1C**)^1^. Western blot analysis confirmed that ARID1B protein levels were significantly reduced in day 6 *Arid1b* KO cells compared with WT cells (**Fig. 1D**).

**Figure 1.**
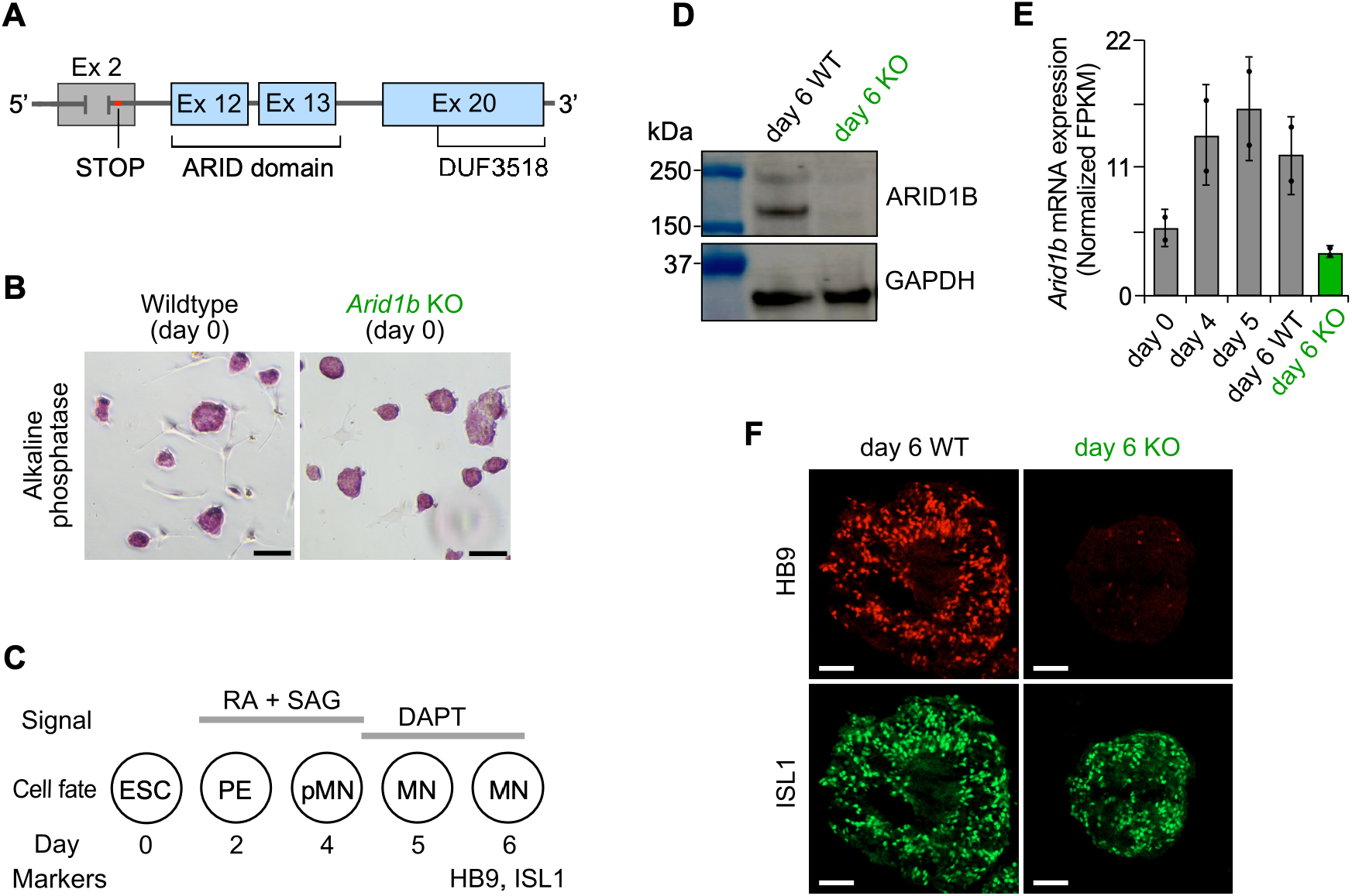
*Arid1b* knockout cells, generated via CRISPR/Cas9, exhibit reduced expression of motor neuron markers. **A.** Schematic representation of the homozygous frameshift mutation in exon 2 of the mouse *Arid1b* gene, resulting in a premature stop codon. **B.** Day 0 mouse embryonic stem (ES) cells stained for alkaline phosphatase activity as a marker of pluripotency. Wild-type (WT) and *Arid1b* KO ES cells exhibited comparable purple staining, indicating similar alkaline phosphatase activity. Scale bars represent 200 µm. **C.** Overview of directed differentiation of mouse ES cells into postmitotic spinal motor neurons (MNs). Differentiating cells progress through primitive ectoderm (PE) on day 2, motor neuron progenitors (pMNs) on day 4, nascent MNs on day 5, and maturing MNs on day 6. RA, retinoic acid; SAG, Sonic Hedgehog pathway agonist; DAPT, γ-secretase inhibitor. **D.** Western blot analysis of ARID1B and GAPDH (loading control) in day 6 WT and *Arid1b* KO cells. Full-length ARID1B and a major isoform were detected at approximately 238 and 191 kDa (kilodaltons), respectively. Unprocessed immunoblot images from two biological replicates from biologically independent cell differentiations (*n* = 2) were shown in Supplementary Data 1. **E.** *Arid1b* mRNA expression measured by RNA-seq on days 0, 4, 5, and 6 of differentiation in WT cells and on day 6 in *Arid1b* KO cells (Supplementary Data 2). FPKM (fragments per kilobase of transcript per million mapped reads) values from two biological replicates derived from independent cell differentiations were median-normalized and averaged (Supplementary Data 3, 4). Data points represent median-normalized FPKM values from individual biological replicates (*n =* 2); error bars represent the standard deviation. **F.** Immunostaining of differentiated embryoid bodies (three-dimensional cell aggregates) for the motor neuron-specific markers HB9 (MNX1) and ISL1 in day 6 WT and *Arid1b* KO cells. Scale bars represent 50 µm.

To characterize gene expression during motor neuron differentiation, we performed RNA sequencing (RNA-seq) of WT and *Arid1b* KO cells at different stages of differentiation. We observed that *Arid1b* expression increased in day 4 WT neural progenitors compared with day 0 WT ES cells and remained elevated through day 6 in WT motor neurons but was significantly reduced in day 6 *Arid1b* KO cells (**Fig. 1E**). Using the day 0 RNA-seq datasets, we also evaluated whether *Arid1b* KO alters genome-wide gene expression in undifferentiated ES cells. Differential gene expression analysis revealed no significant changes in 99.9% of all annotated mRNA genes (19,275 of 19,290) (Supplementary Fig. 1A). Additionally, Gene Ontology analysis showed that genes enriched in day 0 *Arid1b* KO ES cells relative to day 6 motor neurons were associated with ES cell-related processes, similar to the enrichment observed in day 0 WT ES cells (Supplementary Fig. 1B).

We also investigated whether *Arid1b* KO affects the differentiation of ES cells into day 4 neural progenitors. Differential gene expression analysis revealed no significant changes between day 4 WT and *Arid1b* KO cells for more than 99.9% of all annotated mRNA genes (19,287 of 19,290) (Supplementary Fig. 1C). Only three mRNA genes, including *Arid1b*, were significantly differentially expressed. Together with the alkaline phosphatase staining (**Fig. 1B**), these results demonstrate that *Arid1b* KO does not substantially impair ES cell pluripotency or early differentiation into neural progenitors.

Next, to examine whether *Arid1b* KO affects the expression of motor neuron markers, we performed immunostaining for the postmitotic motor neuron markers ISL1 and HB9 in day 6 WT and *Arid1b* KO cells. We observed reduced ISL1 and HB9 immunostaining in day 6 *Arid1b* KO cells compared with WT cells (**Fig. 1F**). Furthermore, mRNA levels of key motor neuron genes (*Hb9*, *Isl1*, and *Lhx3*)*^2C,^*^27^ were significantly reduced in day 6 *Arid1b* KO cells compared with WT cells (Supplementary Fig. 2A). Expression of immature neuronal genes (*Nrp1, Robo1* and *Dcx*)^28–30^ and mature neuronal genes (*Dlg4, Map6,* and *Mapt*)^31–33^ was also significantly reduced in day 6 *Arid1b* KO cells compared with day 6 WT motor neurons (Supplementary Fig. 2B). Overall, these results demonstrate that ARID1B is required for the normal expression of motor neuron and other neuronal genes during motor neuron differentiation.

### ARID1B is required for neuronal gene activation and cell-cycle gene repression

To examine whether loss of ARID1B affects global gene expression, we performed RNA-seq in day 6 WT motor neurons and *Arid1b* KO cells. We found that *Arid1b* KO resulted in significant differential expression of 12.6% of all annotated mRNA genes (2,434 of 19,290) on day 6 of motor neuron differentiation (**Fig. 2A**, left). We categorized these differentially expressed genes into three groups: *i*) highly downregulated (915 genes), *ii*) highly upregulated (843 genes), and *iii*) weakly upregulated (676 genes) in day 6 *Arid1b* KO cells compared with day 6 WT motor neurons. The most highly downregulated genes included mature neuronal and motor neuron-specific markers, such as *Hb9, Lhx3,* and *Hox5a* (**Fig. 2B**). In contrast, the most highly upregulated genes included early developmental genes such as *Lrp2*, which encodes a Sonic Hedgehog receptor^34^, and *Gprc5c*, which encodes a G protein-coupled receptor^35^. Notably, *Apoe*, a genetic risk factor for Alzheimer’s disease^36^, was also highly upregulated in day 6 *Arid1b* KO cells. Additionally, weakly upregulated genes included key mitotic cell-cycle regulators such as *Cdk1, Cdk6,* and *Cdca2*^37,38^. Furthermore, Gene Ontology analysis revealed that highly downregulated genes (Group *i*) were significantly enriched for mature neuronal functions, including chemical synaptic signaling, synaptic transmission, and dendrite organization (**Fig. 2C**). In contrast, upregulated genes (Groups *ii* and *iii*) were enriched for developmental and mitotic cell-cycle processes. Together, these results indicate that ARID1B is required for the expression of neuronal gene programs and the repression of early developmental and mitotic cell-cycle gene programs during motor neuron differentiation.

**Figure 2.**
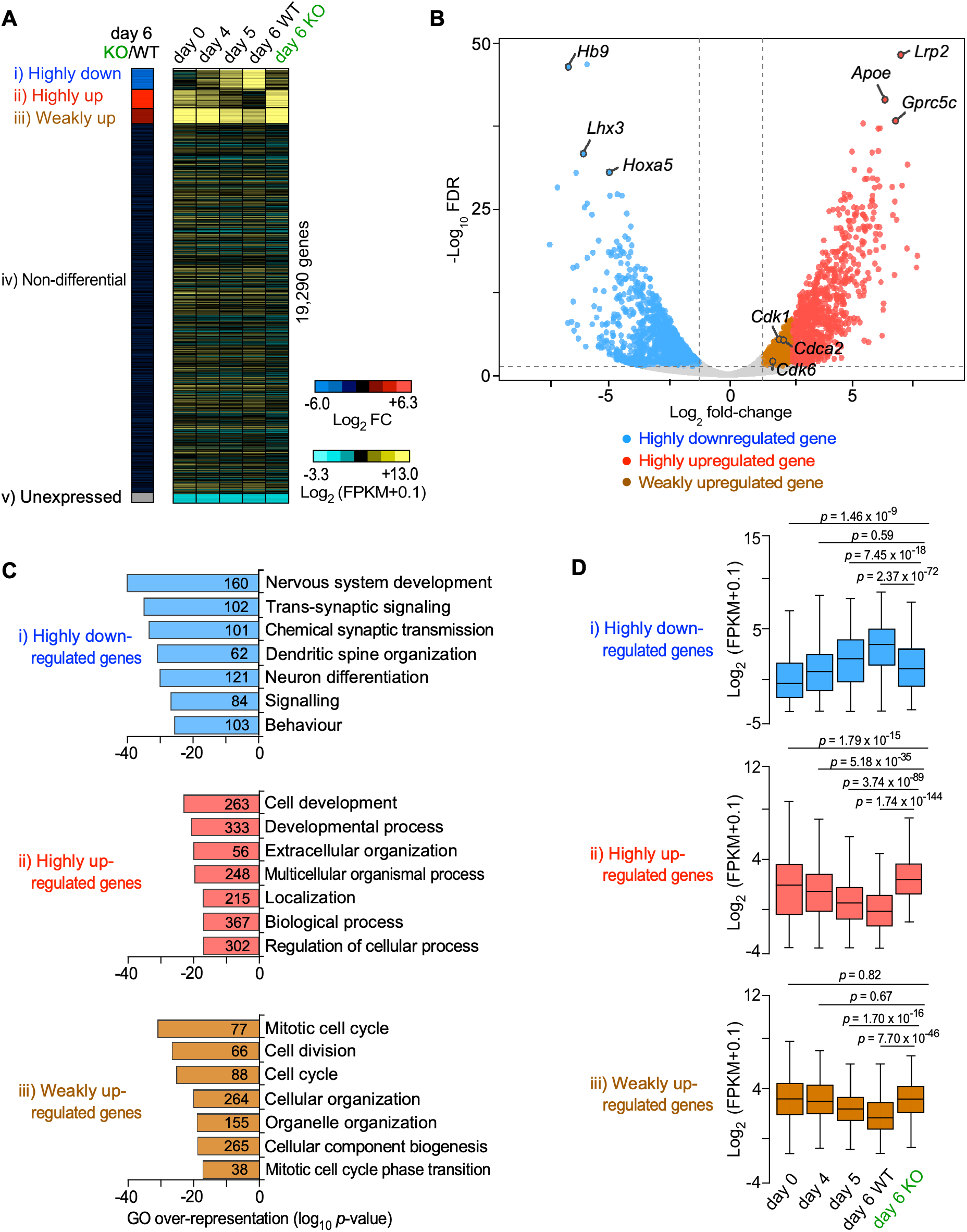
ARID1B is required for neuronal gene activation and repression of mitotic cell-cycle genes. **A.** *Left:* Heatmap of differential gene expression (log_2_ fold-change [FC]) for all 19,290 mRNA genes in day 6 *Arid1b* KO cells compared with day 6 WT motor neurons, measured by RNA-seq using biologically independent cell differentiations (*n* = 2, Supplementary Data 2). Genes were classified into three groups of differentially expressed genes (FC > 2.00, FDR < 0.50 x 10^-3^), identifying 915 highly downregulated (Group *i*, cyan blue), 843 highly upregulated (Group *ii*, light red), and 676 weakly upregulated (Group *iii*, brown) genes. The remaining genes were classified as 16,426 non-differentially expressed (Group *iv*) and 430 unexpressed (Group *v*). *Right:* Heatmap of RNA expression (log_2_ [FPKM+0.1]) of all 19,290 mRNA genes in WT cells on days 0, 4, 5, and 6 and in day 6 *Arid1b* KO cells, measured by RNA-seq. The 19,290 mRNA genes were ordered as displayed on the *left* (Supplementary Data 3). **B.** Volcano plot of all 19,290 mRNA genes showing differential expression between day 6 *Arid1b* KO cells and day 6 WT motor neurons. Differentially expressed gene groups correspond to those defined in panel **A**. **C.** Gene Ontology analysis of the three groups of differentially expressed genes (Groups *i-iii*) defined in panel **A**. The seven most significantly enriched, non-redundant GO terms are shown. Numbers within bars represent the number of genes associated with each GO term (Supplementary Data 5, 6). **D.** Box plots showing RNA expression (log_2_ [FPKM+0.1]) for the three groups of differentially expressed genes (Groups *i-iii*), defined in panel **A**. RNA expression data from biologically independent cell differentiations (*n* = 2) were median-normalized, averaged, and log_2_-transformed. Boxes indicate the interquartile range (IQR), with center lines indicating the median and whiskers extending to 1.5 × IQR (Supplementary Data 4). *P*-values were calculated using two-sided paired *t*-tests. No adjustments for multiple comparisons were performed because each test addressed a distinct, pre-specified hypothesis.

Given that day 6 *Arid1b* KO cells maintained elevated expression of mitotic cell-cycle genes and failed to fully induce postmitotic neuronal genes, we reasoned that day 6 *Arid1b* KO cells might retain transcriptional characteristics of an earlier developmental stage rather than completing motor neuron differentiation by day 6. To test this, we compared the gene expression profile of day 6 *Arid1b* KO cells with those of WT cells at different stages of differentiation (days 0, 4, 5, and 6). We classified all annotated protein-coding mRNA genes into five groups: differentially expressed (Groups *i–iii*), non-differentially expressed (Group *iv*), and unexpressed (Group *v*) genes (**Fig. 2A**, right). Interestingly, expression patterns of differentially expressed genes in day 6 *Arid1b* KO cells most closely resembled those of day 4 WT neural progenitors (**Fig. 2D**). We also confirmed the reproducibility of the RNA-seq data across independent differentiations (Supplementary Fig. 3). Together, these findings indicate that ARID1B is required for the transcriptional transition from proliferating neural progenitors to postmitotic motor neurons but is dispensable for the establishment of neural progenitor identity.

### Differentially expressed gene promoters exhibit distinct histone modification signatures

To examine whether differentially expressed genes exhibit distinct histone modification signatures at their promoters, we performed chromatin immunoprecipitation followed by sequencing (ChIP-seq) for the activating histone marks H3K27ac and H3K4me3 and the repressive histone mark H3K27me3^39,40^. We confirmed that selected genes from each differentially expressed gene group exhibited distinct enrichment patterns for these histone modification marks near their promoters in day 6 motor neurons (**Fig. 3A**, Supplementary Fig. 4A). We next examined genome-wide enrichment of these histone modification marks and found that promoters of highly downregulated genes (Group *i*) were enriched for the activating histone marks H3K27ac and H3K4me3 (**Fig. 3B, C**). Notably, these activating marks were significantly more enriched at promoters of highly downregulated genes (Group *i*) than at those of highly upregulated genes (Group *ii*). In contrast, levels of the repressive histone mark H3K27me3 were significantly higher at promoters of highly upregulated genes (Group *ii*) than at those of highly downregulated genes (Group *i*) (**Fig. 3C, D**). These results demonstrate that the two gene groups exhibit distinct promoter histone modification signatures characterized by differential enrichment of activating and repressive marks in day 6 motor neurons.

**Figure 3.**
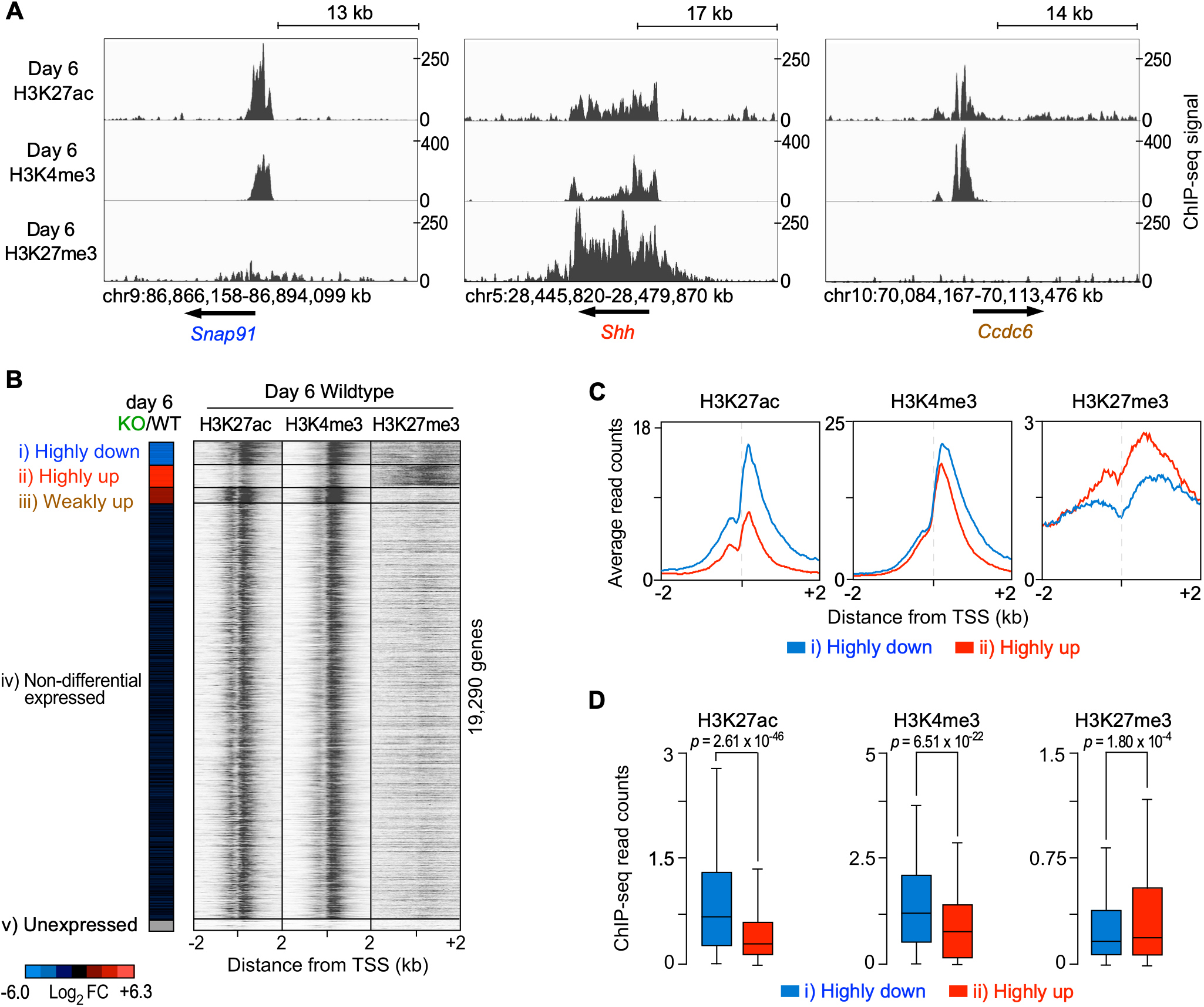
Differentially expressed gene promoters exhibit distinct histone modification signatures in WT motor neurons. **A.** Representative genomic loci for the highly downregulated *SnapS1* (synaptosome-associated protein 91), highly upregulated *Shh* (Sonic Hedgehog), and weakly upregulated *CcdcC* (coiled-coil domain-containing 6) genes, identified in Fig. 2A. H3K27ac, H3K4me3, and H3K27me3 enrichment was measured by ChIP-seq in day 6 WT motor neurons. **B.** *Left*: RNA expression heatmap from Fig. 2A, showing all 19,290 mRNA genes classified into Groups *i–v*. *Right*: Heatmaps of H3K27ac, H3K4me3, and H3K27me3 ChIP-seq signals in day 6 WT motor neurons (Supplementary Data 2). ChIP-seq signals were plotted relative to the transcription start sites (TSSs) of all 19,290 mRNA genes, ordered and grouped according to the RNA expression heatmap on the *left*. **C.** Composite plots of H3K27ac, H3K4me3, and H3K27me3 ChIP-seq signals at promoters of highly downregulated (Group *i*, blue) and highly upregulated (Group *ii*, red) genes defined in panel **B**. Signals represent averages of two biological replicates derived from independent cell differentiations (Supplementary Data 2). **D.** Box plots comparing H3K27ac, H3K4me3, and H3K27me3 ChIP-seq signals at promoters of highly downregulated (Group *i*) and highly upregulated (Group *ii*) genes (Supplementary Data 2). ChIP-seq read counts are the average of two replicates derived from biologically independent cell differentiations (*n =* 2). Boxes indicate the interquartile range (IQR), with center lines indicating the median and whiskers extending to 1.5 × IQR (Supplementary Data 4). *P*-values were calculated using two-sided paired *t*-tests.

This differential enrichment of distinct histone modification marks was further supported by the inverse patterns observed in day 0 WT ES cells (Supplementary Fig. 4B), in which promoters of highly downregulated genes exhibited 1.4-fold greater H3K27me3 enrichment than those of highly upregulated genes (Supplementary Fig. 4C, D). H3K27ac enrichment at these promoters also exhibited an inverse pattern in day 0 ES cells compared with that in day 6 motor neurons. Together, these findings demonstrate that differentially expressed gene groups exhibit distinct and developmentally dynamic promoter histone modification signatures that correspond to their expression patterns during motor neuron differentiation.

### ARID1B and BRG1 co-bind distinct genomic targets in motor neurons

To identify the *cis*-regulatory regions bound by ARID1B-containing BAF complexes, we performed ChIP-seq for ARID1B and the core BAF ATPase BRG1 in day 6 motor neurons. We identified 11,289 ARID1B-bound regions, including regions associated with promoters of key neuronal genes, such as *Irs1* and *Ncam1* (**Fig. 4A**)^41,42^. More than 96.8% of these ARID1B-bound regions (10,928 of 11,289 regions) were co-bound by BRG1 at either distal or proximal *cis*-regulatory regions, corresponding to putative enhancers and promoters, respectively (Supplementary Fig. 5A). These findings are consistent with the incorporation of ARID1B into BRG1-containing BAF complexes, as previously reported^43^. To compare ARID1B/BRG1 co-bound regions with regions lacking ARID1B binding, we further identified BRG1-only bound regions at distal (*n* = 16,955) and proximal (*n* = 6,542) *cis*-regulatory regions. These results indicate that only a subset of BRG1-bound distal *cis*-regulatory regions (6,542 of 21,601) were co-bound by ARID1B (**Fig. 4B**).

**Figure 4.**
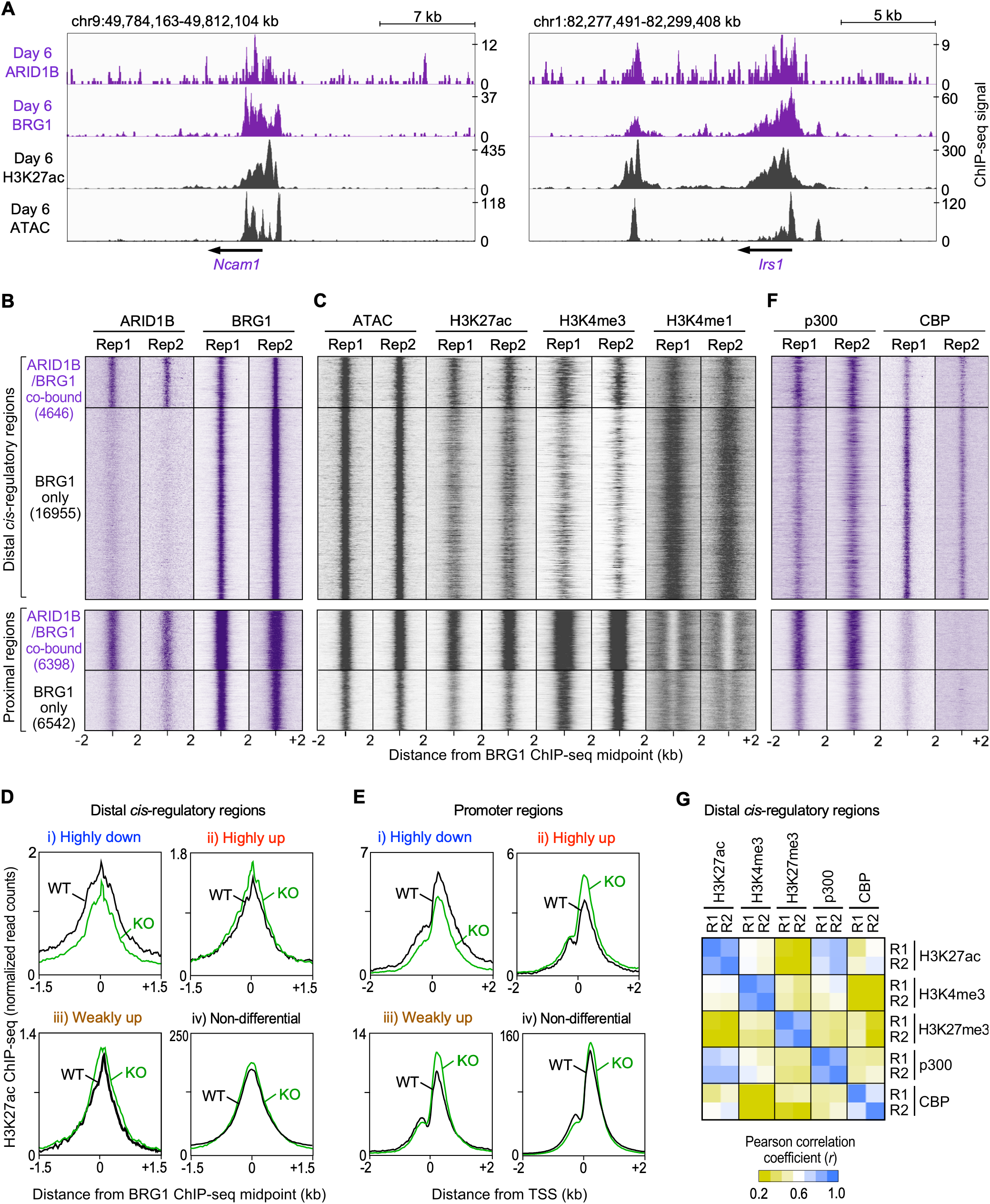
ARID1B and BRG1 co-bind *cis*-regulatory regions associated with distinct H3K27ac enrichment. **A.** Representative genomic loci showing ChIP-seq and ATAC-seq signals at the neuronal cell adhesion molecule *Ncam1* (left) and insulin receptor substrate 1 (*Irs1*) (right) genes, with ARID1B, BRG1, and H3K27ac occupancy in day 6 WT motor neurons. **B.** Heatmaps of ARID1B and BRG1 ChIP-seq signals in day 6 WT motor neurons (Supplementary Data 2). A total of 34,541 regions were classified as ARID1B/BRG1 co-bound (*n* = 4646) or BRG1-only bound (*n* = 16,955) regions at distal *cis*-regulatory regions (*upper* panel) and ARID1B/BRG1 co-bound (*n* = 6398) or BRG1-only bound (*n* = 6542) regions at proximal *cis*-regulatory regions (*lower* panel). Data from two biological replicates derived from independent cell differentiations (replicates 1 and 2) are shown. All signals were plotted relative to day 6 BRG1 ChIP-seq peak midpoints (Supplementary Data 7). **C.** Heatmaps of ATAC-seq signals and H3K27ac, H3K4me3, and H3K4me1 ChIP-seq signals in day 6 WT motor neurons (Supplementary Data 2). Data from two biological replicates derived from independent cell differentiations (replicates 1 and 2) are shown. Data were grouped and ordered as in panel **B**. **D, E.** Composite plots of H3K27ac ChIP-seq signals at ARID1B/BRG1 co-bound distal *cis*-regulatory regions (**D**) and promoters (**E**) associated with four groups of genes (Groups *i–iv*) defined in Fig. 2A. Signals represent averages of two biological replicates derived from independent cell differentiations. **F.** Heatmaps of ChIP-seq of p300 and CBP in day 6 motor neurons (Supplementary Data 2). Data from two biological replicates from independent cell differentiations (replicate 1 and 2) are shown. Data were grouped and ordered as displayed in panel **B**. **G.** Pearson correlation coefficients (*r*) between ChIP-seq samples for histone modifications and acetyltransferases were calculated using log_2_-transformed read counts across 21,601 distal *cis*-regulatory regions defined in panel **B** from two biological replicates derived from independent cell differentiations (R1 and R2). The heatmap shows pairwise correlations among all examined chromatin-associated signals (Supplementary Data 8).

Next, we examined chromatin accessibility and histone modifications at distal *cis*-regulatory regions (termed distal regions) using the assay for transposase-accessible chromatin with sequencing (ATAC-seq) and ChIP-seq. Although both ARID1B/BRG1 co-bound and BRG1-only distal regions showed similar levels of chromatin accessibility, their histone modification profiles differed significantly (**Fig. 4C**, upper panel). ARID1B/BRG1 co-bound distal regions were enriched for the active enhancer-associated histone marks H3K27ac and H3K4me3, whereas BRG1-only bound distal regions were enriched primarily for the poised enhancer-associated histone mark histone H3 lysine 4 monomethylation (H3K4me1) (Supplementary Fig. 5B). These results suggest that ARID1B and BRG1 preferentially co-bind distal *cis*-regulatory regions with active enhancer-associated chromatin signatures, whereas BRG1-only bound regions exhibit H3K4me1-enriched, potentially poised enhancer-associated chromatin signatures.

We further assessed chromatin accessibility and histone modifications at proximal *cis*-regulatory regions (termed proximal regions) (**Fig. 4C**, lower panel). We found that ARID1B/BRG1 co-bound proximal regions were highly accessible, whereas BRG1-only bound proximal regions exhibited significantly lower accessibility (Supplementary Fig. 5C). Furthermore, ARID1B/BRG1 co-bound proximal regions exhibited greater enrichment of the active histone marks H3K27ac and H3K4me3. Genes associated with ARID1B/BRG1 co-bound proximal regions also displayed higher mRNA expression levels than those associated with BRG1-only bound proximal regions (Supplementary Fig. 5D). These results suggest that ARID1B and BRG1 preferentially co-bind highly accessible and transcriptionally active proximal *cis*-regulatory regions in day 6 motor neurons. Taken together, these findings demonstrate that ARID1B-containing BAF complexes bind distinct genomic loci characterized by specific chromatin signatures in day 6 motor neurons.

### ARID1B-associated H3K27ac dynamics correspond to gene expression changes

To investigate whether ARID1B and BRG1 co-bind distal *cis*-regulatory regions associated with differentially expressed genes (**Fig. 3B**), we assigned these regions to their nearest genes. We found that 20.3% of ARID1B/BRG1 co-bound distal regions (943 of 4646) were linked to differentially expressed genes (Groups *i-iii*). We next examined changes in H3K27ac enrichment at these distal regions following *Arid1b* KO (**Fig. 4D**). H3K27ac levels at distal regions linked to highly downregulated genes (Group *i*) were significantly decreased in day 6 *Arid1b* KO cells compared with WT cells (**Fig. 4D**, Supplementary Fig. 6). In contrast, H3K27ac levels at distal regions linked to upregulated genes (Groups *ii-iii*) showed no significant decrease following *Arid1b* KO.

We also mapped day 6 ARID1B and BRG1 ChIP-seq signals relative to the transcription start sites (TSSs) of all annotated protein-coding genes (Supplementary Fig. 7A). Overall, 15.8% of ARID1B/BRG1 co-bound promoters (1009 of 6398 regions) were associated with differentially expressed genes (Groups *i-iii*). The remaining ARID1B/BRG1 co-bound promoters were associated with non-differentially expressed genes (Group *iv*). These genes were robustly and ubiquitously expressed throughout motor neuron differentiation (Supplementary Fig. 7B, C), and their associated Gene Ontology terms were enriched for fundamental cellular and metabolic processes (Supplementary Fig. 7D). These findings suggest that ARID1B-containing BAF complexes also occupy promoters of broadly expressed genes involved in fundamental cellular functions. Consistent with our observations at distal regions, changes in promoter H3K27ac enrichment closely correlated with changes in gene expression following *Arid1b* KO (**Fig. 4E**, Supplementary Fig. 7E). Together, these results demonstrate that H3K27ac dynamics at ARID1B/BRG1 co-bound regions are closely associated with gene expression changes following *Arid1b* KO.

Next, we investigated whether specific histone acetyltransferases (HATs) occupy these H3K27ac-enriched, ARID1B/BRG1 co-bound regions. We therefore performed ChIP-seq for the HATs p300 and CBP in day 6 motor neurons. Notably, p300, but not CBP, was preferentially enriched at ARID1B/BRG1 co-bound regions exhibiting high H3K27ac levels, whereas BRG1-only bound regions with low H3K27ac levels showed lower p300 occupancy (**Fig. 4F**). Pearson correlation analysis further demonstrated that H3K27ac occupancy correlated more strongly with p300 occupancy (*r* = 0.81) than with CBP (*r* = 0.53) at distal regions bound by ARID1B or BRG1 (**Fig. 4G**). Moreover, p300 occupancy correlated more strongly with H3K27ac (*r* = 0.82) than with H3K4me3 (*r* = 0.59) and H3K27me3 (*r* = −0.07). We also confirmed the reproducibility of the ChIP-seq and ATAC-seq data across independent differentiations (Supplementary Fig. 8). Together, our findings suggest that p300 is associated with H3K27ac enrichment at ARID1B/BRG1 co-bound enhancers and promoters, supporting a model in which ARID1B-containing BAF complexes coordinate local histone acetylation and gene expression during motor neuron differentiation.

### ARID1B/BRG1 co-bound enhancers and promoters are enriched for distinct TF motifs

We next investigated whether ARID1B/BRG1 co-bound distal regions were enriched for specific TF motifs. Motif enrichment analysis revealed that these co-bound distal regions were enriched for CTCF, RFX family, and SOX family TF motifs (**Fig. 5A**). Interestingly, individual motif occurrence analysis^44^ showed that ARID1B/BRG1 co-bound distal regions associated with highly upregulated and non-differentially expressed gene groups (Groups *ii, iv*) were most strongly enriched for the CTCF motif (**Fig. 5B**). To validate these findings, we examined CTCF occupancy in day 6 motor neurons. Consistent with the motif enrichment analysis, ARID1B/BRG1 co-bound distal regions exhibited 1.6-fold greater CTCF occupancy than BRG1-only distal regions (**Fig. 5C**). Conversely, ARID1B/BRG1 co-bound distal regions associated with highly downregulated neuronal genes (Group *i*) were enriched for RFX family TF motifs, which have been implicated in neuronal development and cell proliferation^45,46^. These results suggest that ARID1B-containing BAF complexes occupy distal *cis*-regulatory regions characterized by distinct TF motif and binding signatures associated with different gene expression programs.

**Figure 5.**
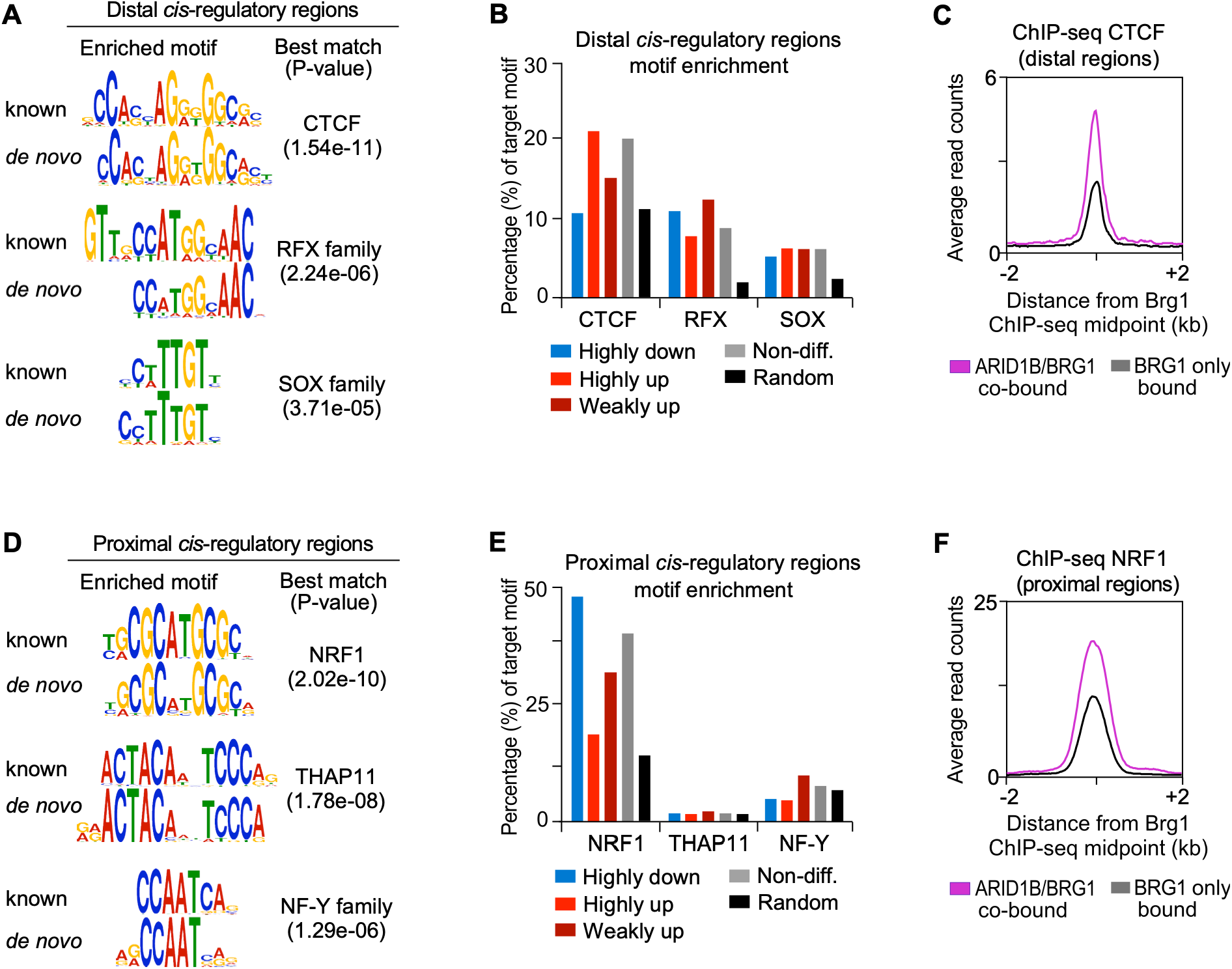
ARID1B/BRG1 co-bound proximal and distal regions exhibit distinct transcription factor binding signatures. **A, D.** Motif enrichment analysis of 4646 ARID1B/BRG1 co-bound distal *cis-*regulatory regions (**A**) and 6398 ARID1B/BRG1 co-bound proximal *cis*-regulatory regions (**D**), defined in Fig. 4B. The *de novo* DNA motifs represent the three most significantly enriched TF-associated motifs identified within 100 bp of ATAC-seq peak midpoints. **B, E.** Frequency of occurrence of the TF-associated motifs identified in panels **A** and **D** across genomic regions associated with gene Groups *i–iv*, defined in Fig. 2B, and randomly selected genomic regions (*n* = 300) (Supplementary Data 9). **C, F.** Composite plots of CTCF ChIP-seq signals at the 4646 ARID1B/BRG1 co-bound distal regions and 16,965 BRG1-only bound distal regions (**C**) and NRF1 ChIP-seq signals at 6398 ARID1B/BRG1 co-bound proximal regions and 6542 BRG1-only bound proximal regions (**F**) in day 6 WT motor neurons.

We also examined TF motifs at ARID1B/BRG1 co-bound proximal regions. Motif enrichment analysis revealed enrichment of NRF1, THAP11, and NF-Y family TF motifs (**Fig. 5D**). We found that ARID1B/BRG1 co-bound proximal regions across all differentially expressed gene groups (Groups *i-iii*) were most strongly enriched for the NRF1 motif (**Fig. 5E**). Furthermore, ARID1B/BRG1 co-bound proximal regions exhibited greater NRF1 occupancy than those bound by BRG1 only (**Fig. 5F**). Given that NRF1 has been implicated in transcriptional activation during the transition from neural progenitors to postmitotic motor neurons^47^, the enrichment of NRF1 motifs and occupancy at ARID1B/BRG1 co-bound promoters identifies NRF1 as a candidate TF that may facilitate recruitment of ARID1B-containing BAF complexes to these regions during motor neuron differentiation. Together, these findings suggest that ARID1B-containing BAF complexes occupy distal and proximal *cis*-regulatory regions characterized by distinct TF motif and occupancy signatures, potentially enabling TF-specific regulation of gene expression during motor neuron differentiation.

### ARID1B promotes cell-cycle exit during the transition from progenitors to motor neurons

Given that day 6 *Arid1b* KO cells exhibit increased expression of mitotic cell-cycle genes (**Fig. 2**), we investigated whether *Arid1b* is required for cell-cycle regulation during motor neuron differentiation. To assess cell proliferation in WT and *Arid1b* KO neural progenitors, we labeled cells with bromodeoxyuridine (BrdU) on day 4 and performed BrdU immunostaining on day 5 (**Fig. 6A**). We also examined whether *Arid1b* is required for the reduction in cell proliferation during differentiation by labeling cells with BrdU on day 5 and performing BrdU immunostaining on day 6 (**Fig. 6B**). Notably, we observed a 2.8-fold decrease in the proportion of BrdU-positive WT cells between the day 4 and day 5 BrdU-labeling experiments, whereas the proportion of BrdU-positive *Arid1b* KO cells remained unchanged (**Fig. 6C**). Consequently, the proportion of BrdU-positive *Arid1b* KO cells was significantly higher than that of WT cells following the day 5 BrdU-labeling experiments. These results indicate that the proportion of proliferating WT cells decreases between days 4 and 6, whereas *Arid1b* KO cells maintain a higher level of proliferation during this period. Furthermore, we confirmed that *Arid1b* KO cells fail to repress mitotic cell-cycle genes (*Cdk1, Cdk2,* and *Cdca2*)^37,38,48^ beyond day 4 of differentiation (Supplementary Fig. 9). Together, these results suggest that ARID1B is required for the repression of cell-cycle genes and promotes the transition from proliferating neural progenitors to postmitotic motor neurons by facilitating cell-cycle exit.

**Figure 6.**
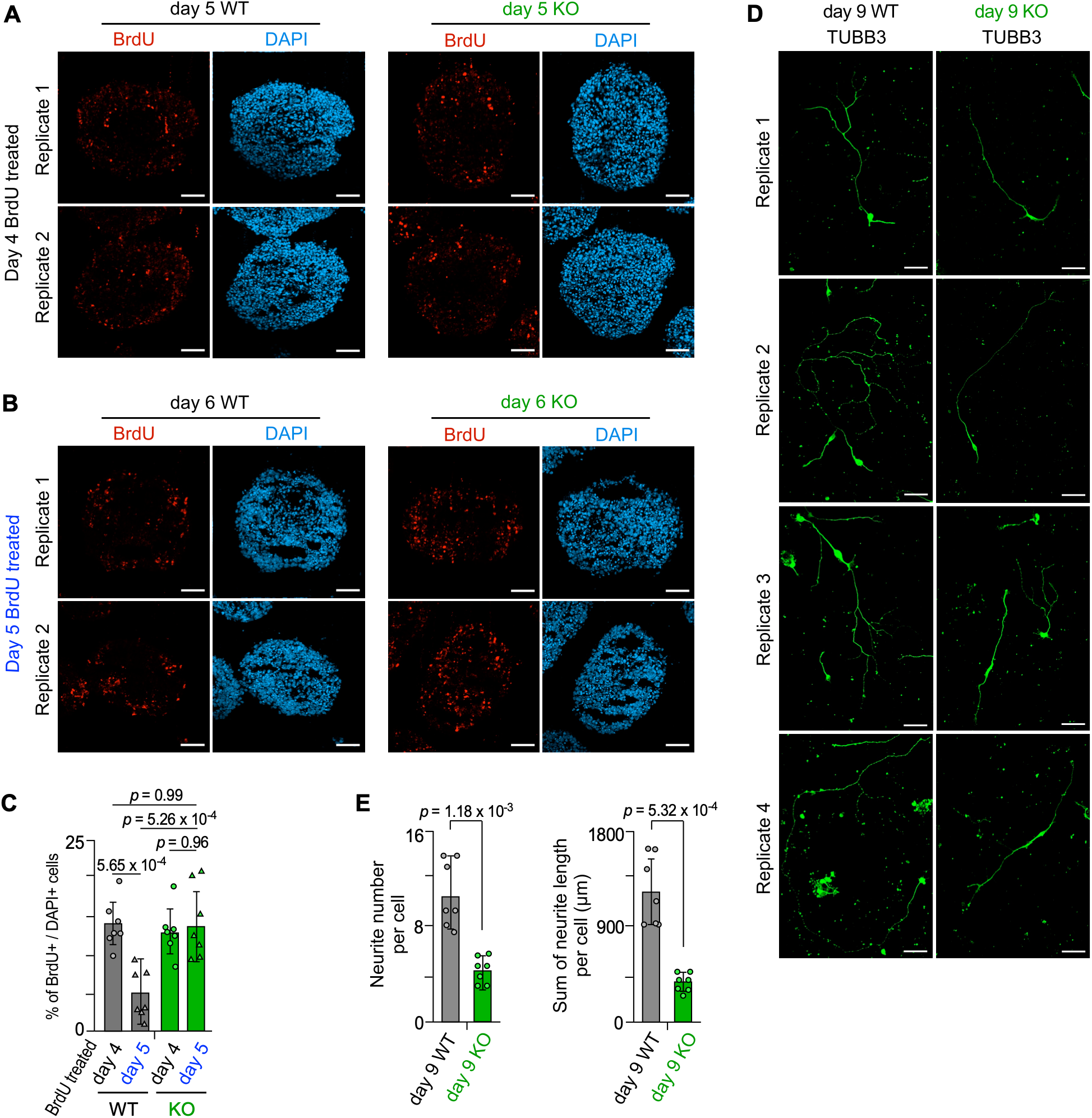
ARID1B promotes cell-cycle exit during the transition from neural progenitors to postmitotic motor neurons. **A.** WT and *Arid1b* KO embryoid bodies were labeled with BrdU on day 4 and collected on day 5 for immunostaining for BrdU and staining with the nuclear dye DAPI (4’,6-diamidino-2-phenylindole). Scale bars represent 50 µm. Two representative images from two biological replicates derived from independent cell differentiations are shown. **B.** BrdU labeling and immunostaining were performed as described in panel **A**, except that embryoid bodies were labeled with BrdU on day 5 and collected on day 6. **C.** The percentage of BrdU-positive cells among DAPI-positive cells was quantified in WT and *Arid1b* KO embryoid bodies following BrdU labeling on day 4 or day 5. Quantification includes data from biologically independent cell differentiations (*n* = 2). Data points represent values from individual embryoid bodies (*n =* 7, Supplementary Data 10). Error bars represent the standard deviation. *P*-values were calculated using one-way ANOVA followed by Tukey’s honestly significant difference (HSD) test (Supplementary Data 11). **D.** Immunostaining of day 9 WT and *Arid1b* KO cells for the neuronal marker TUBB3. Day 6 embryoid bodies were dissociated into single cells and cultured for an additional three days on laminin-coated surfaces. Scale bars represent 50 µm. Four representative images from two biological replicates derived from independent cell differentiations are shown. **E.** *Left:* Number of TUBB3-positive neurites per day 9 WT or *Arid1b* KO cell. *Right*: Total neurite length (µm) per day 9 WT or *Arid1b* KO cell. Quantification includes data from biologically independent cell differentiations (*n* = 2). Data points represent values from individual cells (*n =* 7, Supplementary Data 12). Error bars represent the standard deviation. *P*-values were calculated using two-sided paired *t*-tests.

Finally, we investigated whether *Arid1b* KO impairs neurite outgrowth in differentiated neurons. To address this, we extended the differentiation period by three days to generate day 9 maturing motor neurons and visualized their morphology by immunostaining for TUBB3, a neuronal marker that labels cell bodies and neurites^49^. We observed a marked reduction in neurite complexity of day 9 *Arid1b* KO cells compared with day 9 WT motor neurons (**Fig. 6D**). Specifically, *Arid1b* KO cells exhibited significant reductions in both neurite number and length per neuron compared with WT motor neurons (**Fig. 6E**). These results suggest that ARID1B is required for proper neurite outgrowth during motor neuron maturation. Taken together, our findings demonstrate that ARID1B promotes motor neuron differentiation, cell-cycle exit, and neurite outgrowth, consistent with its requirement for the repression of early developmental and mitotic gene expression programs and the activation of mature neuronal gene expression programs.

## Discussion

ARID1B is temporally incorporated into npBAF and nBAF complexes, which regulate neuronal differentiation by controlling the expression of lineage-commitment genes through chromatin remodeling^14,15^. *Arid1b* mutations have been linked to a variety of central nervous system disorders in humans, highlighting the importance of ARID1B in neurodevelopment^16^. However, the effects of *Arid1b* mutations on neuronal cell fate acquisition across specific neuronal lineages remain poorly understood. Here, we investigated the effects of *Arid1b* mutation on spinal motor neuron specification, given the reported motor system dysfunction in patients with *Arid1b* mutations and *Arid1b* haploinsufficient mice^17,18^.

We found that ARID1B is required for the repression of mitotic cell-cycle genes and promotes neural progenitor cell-cycle exit during motor neuron differentiation (**Fig. 2**). Previous studies have similarly implicated ARID1B in regulating cell-cycle dynamics during cortical neurogenesis^50,25^. However, the effects of ARID1B deficiency on neural progenitor proliferation vary across neuronal populations. The integration of ARID1B into nBAF is essential for neuroectoderm lineage commitment, as *Arid1b* haploinsufficiency leads to persistent expression of pluripotency factors and associated gene networks^14^. In an analysis of inhibitory neuron differentiation in *Arid1b* haploinsufficient mouse brains, Jung *et al*.^19^ demonstrated a significant reduction in postmitotic GABAergic neurons compared with WT controls. This deficit was attributed to decreased ventral progenitor proliferation and elevated apoptosis within the medial ganglionic eminence (MGE)^19^. Building on these findings, Moffat *et al.*^50^ showed that impaired proliferation resulted from reduced cell-cycle speed and compromised cell-cycle re-entry following ARID1B loss. Comparison with dorsal excitatory progenitors in the ventricular zone revealed distinct proliferative outcomes, highlighting a cell type-specific role for ARID1B in neural progenitor expansion^50^. In contrast, our findings demonstrate that *Arid1b* KO motor neuron progenitors maintain elevated proliferation during the transition to postmitotic motor neurons, suggesting that ARID1B regulates cell-cycle dynamics in a lineage- and developmental-stage-specific manner.

We further found that ARID1B is required for normal expression of mature neuronal genes and proper neurite outgrowth during motor neuron differentiation (**Fig. 6**). Previous studies have shown that *Arid1b* mutations impair neurite outgrowth and synaptic morphology in cortical neurons^19–21^. Our findings extend these observations to spinal motor neurons and integrate this morphological phenotype with genome-wide analyses of ARID1B-containing BAF complexes. Proper motor system function depends on the specification, maturation, and connectivity of motor neurons^22^. We show that *Arid1b* KO impairs motor neuron differentiation and neurite development, providing a potential cellular mechanism contributing to the motor dysfunction associated with *Arid1b* haploinsufficiency in humans and mice^17,18^. How these cellular defects affect motor neuron connectivity and circuit function remains unknown. Future studies using *in vivo* models will be necessary to determine whether alterations in motor neuron number, maturation, and morphology contribute to impaired motor circuit assembly and function following ARID1B loss.

Cell type-specific subunits of the BAF complex are thought to modulate genomic targeting and occupancy at *cis*-regulatory regions^8,9,11,12^. Thus, characterizing the genomic occupancy of cell type-specific BAF complexes is important for understanding their roles during differentiation. In this study, we performed genome-wide mapping of ARID1B and BRG1 occupancy in WT motor neurons and characterized the associated chromatin signatures. We found that ARID1B and BRG1 co-bind gene promoters and distal *cis*-regulatory regions with chromatin signatures distinct from those of regions bound by BRG1 alone (**Fig. 4**). Specifically, ARID1B and BRG1 co-bind highly accessible and active gene promoters that exhibit greater enrichment of H3K27ac and H3K4me3 than promoters occupied by BRG1 alone. Additionally, ARID1B and BRG1 co-bound distal *cis*-regulatory regions were enriched for active chromatin signatures, including H3K27ac, whereas BRG1-only distal regions were enriched for H3K4me1 and exhibited lower H3K27ac enrichment, consistent with a poised enhancer-like chromatin state. These results suggest that incorporation of cell type-specific subunits may contribute to the distinct genomic occupancy and chromatin states of mammalian BAF complexes.

Many studies have highlighted functional interactions between the BAF complex and cell type-specific TFs that facilitate recruitment to the *cis*-regulatory regions of lineage-commitment genes^51^. ARID1B contains a conserved ARID domain that facilitates DNA binding^25^. However, ARID1B binds DNA without apparent sequence specificity^43,52^. Therefore, we propose that ARID1B-containing BAF complexes are recruited to target *cis*-regulatory elements through interactions with specific TFs. Our TF motif and occupancy analyses identified NRF1 as a candidate TF associated with ARID1B/BRG1 co-bound promoters in WT motor neurons (**Fig. 5**).

Consistent with the established pioneer TF activity of NRF1 during mouse ES cell differentiation^53^, we propose that NRF1 may establish or maintain chromatin accessibility at target promoters, thereby facilitating recruitment of ARID1B-containing BAF complexes. Because ARID1B lacks intrinsic enzymatic activity to directly modify histones, we propose a model in which ARID1B-containing BAF complexes facilitate interactions between regulatory chromatin and histone-modifying enzymes.

Supporting this model, we showed that the histone acetyltransferase p300, but not CBP, preferentially occupies H3K27ac-enriched ARID1B/BRG1 co-bound *cis*-regulatory regions in WT motor neurons. These findings complement recent work demonstrating that *Arid1b* haploinsufficiency reduces CBP recruitment and impairs histone acetylation in the mouse cortex^19^, suggesting that ARID1B-containing BAF complexes may cooperate with distinct histone acetyltransferases in a cell type-specific manner. Similarly, Liu *et al.*^47^ reported that NRF1 recruits an AUTS2-ncPRC1.3-p300 complex to motor neuron promoters, promoting H3K27ac deposition and activating gene expression programs essential for motor neuron specification. Together, these findings support a model in which NRF1, ARID1B-containing BAF complexes, and p300 cooperate at neuronal regulatory regions to establish transcriptionally permissive chromatin during motor neuron differentiation. Future studies should determine the molecular interactions among these factors and clarify how ARID1B-containing BAF complexes coordinate local histone acetylation at *cis*-regulatory elements to fine-tune gene expression during cell-fate commitment.

Furthermore, our TF motif and occupancy analyses revealed that distal *cis*-regulatory regions co-bound by ARID1B and BRG1 were significantly enriched for CTCF and RFX family motifs, with preferential CTCF occupancy at ARID1B/BRG1 co-bound distal regions (**Fig. 5**). Thus, we further propose that ARID1B may function beyond direct transcriptional control at *cis*-regulatory regions associated with non-differentially expressed genes. Mammalian BAF complexes are known to localize to *cis*-regulatory regions and regulate non-transcriptional processes, including higher-order chromatin organization, genomic stability, and DNA repair^54^. For instance, Euskirchen *et al.*^55^ demonstrated that approximately 17% of BAF binding sites in HeLa cells overlap with CTCF-bound regions, indicating that BAF complexes localize not only to canonical promoters and enhancers but also to architectural genomic boundaries. Consistent with these findings, our motif enrichment and ChIP-seq analyses revealed significant CTCF motif enrichment and preferential CTCF occupancy at ARID1B/BRG1 co-bound distal *cis-*regulatory regions, including those associated with non-differentially expressed genes (**Fig. 5**). Euskirchen *et al.*^55^ further established that BAF complexes assemble into diverse combinatorial structures, supporting their versatile transcriptional and non-transcriptional roles. Together, our findings and previous studies raise the possibility that ARID1B-containing BAF complexes contribute to higher-order chromatin organization in motor neurons, potentially through distinct subunit compositions. Future work should determine whether ARID1B-containing BAF complexes directly regulate chromatin architecture and how such functions are coordinated with their transcriptional roles during cell-fate commitment.

Taken together, our findings demonstrate that *Arid1b* mutation disrupts spinal motor neuron specification. We establish that ARID1B is essential for the balanced expression of early developmental, mitotic cell-cycle, and mature neuronal gene networks during motor neuron differentiation. ARID1B-containing BAF complexes occupy distinct *cis*-regulatory elements, where ARID1B loss is associated with altered H3K27ac enrichment and gene expression. Functionally, ARID1B is required for cell-cycle exit, efficient motor neuron differentiation, and proper neurite outgrowth. Future studies will be critical for identifying cofactors, including specific histone-modifying enzymes and cell type-specific TFs, that partner with ARID1B-containing BAF complexes to regulate chromatin and transcription. Overall, this study provides insight into how ARID1B-containing BAF complexes coordinate transcriptional and epigenetic programs during motor neuron differentiation, advancing our understanding of how specialized BAF subunits contribute to neuronal cell identity and how their disruption may contribute to neurodevelopmental disorders.

## Methods

### Cell culture

The embryonic stem (ES) cell line originated from the male embryo of the inbred mouse strain 129/Ola ES-E14. Mouse ES cells were cultured and differentiated into spinal motor neurons as described previously^1^. Briefly, ES cells were plated on 0.1% gelatin-coated plates in EmbryoMax DMEM (Millipore, SLM220) supplemented with 10% ES cell-grade fetal bovine serum (Invitrogen, 12483020), L-glutamine (Gibco, 25030081), 0.1 mM beta-mercaptoethanol, and leukemia inhibitory factor (Millipore, ESG1107). After two days, we trypsinized the ES cells (Thermo, 25300054) and seeded them at 7.5 x 10^4^ cells per mL in 1:1 Advanced DMEM/F12 (Thermo, 12634010) and Neurobasal Medium (Gibco, 21103049) supplemented with 10% Knockout Serum Replacement (Thermo, 10828028), 2 mM L-glutamine (Gibco, 25030081), and 0.1 mM beta-mercaptoethanol to initiate the formation of embryoid bodies (day 0). The media was replaced on days 2 and 5 of differentiation. Patterning of embryoid bodies was induced by supplementing the media on day 2 with 1 μM all-trans retinoic acid (Sigma, R2625) and 0.25 μM Smoothened agonist (SMO) of hedgehog signaling (Selleckchem, S7779) and on day 4 with 5 μM DAPT (Selleckchem, S2215).

For neuronal culture, embryoid bodies of differentiated motor neurons were collected on day 6, trypsinized into single cells, and plated for attachment on coverslips coated with 0.001% poly-l-ornithine (Sigma, A-004-M) and 1% Matrigel (VWR, 356234) in 1:1 Advanced DMEM/F12 (Thermo, 12634010) and Neurobasal Medium (Gibco, 21103049) supplemented with 2% B-27 (Gibco, A3582801), Horse Serum (Gibco, 16050130), 2 mM L-glutamine (Gibco, 25030081), 0.1 mM beta-mercaptoethanol, and Pen/Strep (Thermo, 15140122). The media was replaced on day 8. Cells attached on coverslips were collected to day 9 for immunofluorescence.

### Generation of *Arid1b* knockout ES cell line

The RNA-guided Cas9 nuclease from the microbial Clustered Regularly Interspaced Short Palindromic Repeats (CRISPR/Cas9) system was used to delete 56 bp of *Arid1b* in ESCs. The CRISPR/Cas9 genome editing was performed as described in Ran *et al.*^56^, with some modifications. Briefly, guide RNAs (gRNAs) were designed using available software, including CRISPOR (http://crispor.tefor.net/crispor.py) and CHOPCHOP (https://chopchop.cbu.uib.no/). A pair of oligonucleotides encoding the 23 bp guide sequences containing the 5’-NGG PAM (protospacer adjacent motif) sequence (underlined) to target the left and right sides of the intended deletion. gRNA sequences target the 56 bp region of the *Arid1b* (chromosome 17): 5’ GGAGGCTCCTACGGTCCCCCAGG 3’ (forward strand), Reverse 5’ CGGGTACTGCAAGCCTCCCAGGG 3’ (reverse strand). The 23 bp guide sequences were synthesized from Integrated DNA Technologies, Inc. (Coralville, IA, USA). To synthesize gRNAs, we cloned 23 bp genomic sites of the form 5’-N20NGG-3’ (N=A, T, G, or C) into the U6 target gRNA expression vector (pgRNA vector) containing the U6 promoter (Addgene, 41824) following the available protocol (https://www.addgene.org/static/cms/files/hCRISPR_gRNA_Synthesis.pdf).

To generate the *Arid1b* KO ES cell line, ES cells were seeded and incubated for 48 hours on mouse embryonic fibroblast cells (EmbryoMax, PMEF-N) to maintain stem cell pluripotency. Cells were transiently transfected with a pair of pgRNA vectors and a pCas9-GFP plasmid (Addgene, 44719) containing a CAGGS promoter with Cas9 DNA endonuclease and a GFP reporter using the Mouse P3 Primary Cell Nucleofector Kit (Lonza, VPG-1004) and a Lonza Amaxa 4D-Nucleofector, according to the manufacturer’s instructions. Transfected ES cells were seeded and grown for 24-48 hours after transfection before we trypsinized the ES cells (Thermo, 25300054) and selected GFP-expressing ES cells by fluorescence-activated cell sorting (FACS). Approximately 2,000 sorted cells were plated on a dish with mouse feeder cells. ESCs formed clones 4-6 days after FACS sorting and plating. A single colony was picked, followed by genotyping using PCR primers around genomic regions of intended deletion. The ES cell clone containing the desired deletion was subcultured onto mouse feeder cells and used for subsequent motor neuron differentiation.

### Alkaline phosphatase assay

WT and *Arid1b* KO ES cells were plated on 0.1% gelatin-coated plates in EmbryoMax DMEM (Millipore, SLM220) supplemented with 10% ES cell-grade fetal bovine serum (Invitrogen, 12483020), L-glutamine (Gibco, 25030081), 0.1 mM beta-mercaptoethanol, and leukemia inhibitory factor (Millipore, ESG1107). After two days, staining was performed using an Alkaline Phosphatase Staining Kit (Abcam, ab242286). ES cell colonies were imaged at 10X magnification under a light microscope.

### Western Immunoblotting

Immunoblotting was performed on total cell lysates collected from approximately 50 million cells. Lysates were diluted in 4X Laemmli Sample Buffer (Bio-Rad, 1610747), heated at 95 degrees for 10 minutes, loaded onto Mini-Protean TGX Precast Gels (Bio-Rad, 4561085), and run at 140 V for 1 hour. Transfer was performed onto 0.45 mm PVDF membranes (Sigma, IPVH07850) at 100 V for 3 hours. Gel electrophoresis and transfer were performed using the Mini-PROTEAN® Tetra Cell (Bio-Rad, 1658033). The membranes were incubated with primary antibodies overnight at 4 degrees. Primary antibodies used in this study are anti-GAPDH (1:5000 used, Cell Signalling, 14C10, lot# 14) and anti-ARID1B (1:500 used, Abcam, ab57461, lot# 1144777-1). For chemiluminescence detection of primary antibodies, membranes were incubated for 2 hours with HRP-conjugated goat anti-mouse secondary antibodies (1:10,000 used, Santa Cruz, SC-2005), developed with SuperSignal West Pico PLUS Chemiluminescent Substrate (Thermo, 34580), and imaged with the iBright FL1500 Imaging System.

### BrdU assay

Motor neuron culture media were supplemented with 10 uM bromodeoxyuridine (BrdU) diluted in dimethyl sulfoxide (DMSO) on days 4 and 5 of differentiation for 5 hours. As a control, motor neuron culture medium was supplemented with DMSO only on day 5 of differentiation for 5 hours. After 5 hours, the media was removed, and the embryoid bodies were washed three times using PBS. Embryoid bodies were cultured for an additional 24 hours in fresh motor neuron culture media prior to being collected for immunofluorescence detection of BrdU.

### Immunofluorescence

Day 5 and day 6 embryoid bodies containing differentiated motor neurons were collected from culture media, washed once with ice-cold PBS, and fixed with 4% paraformaldehyde in PBS for 5 minutes at room temperature. Fixed embryoid bodies were washed twice with ice-cold PBS, embedded in Tissue-Tek OCT (VWR, 25608-930), and cryosectioned. Slides were washed once with ice-cold PBS, permeabilized with 0.35% Triton-X for 10 minutes, and blocked for 30 minutes at room temperature. The slides were then incubated with primary antibodies at 4 degrees overnight. Primary antibodies used in this study are anti-BrdU (1ug/mL used, Invitrogen, MA3-071, lot# 3211890), anti-ISL1 (1ug/mL used, RCD Systems, AF1837-SP, lot# JWV0523021), and anti-HB9 (1:1000 used, DSHB, 81.5C10-s, lot# NA). Slides were washed three times with ice-cold PBS and incubated with anti-goat (1:1000 used, Thermo, A11055) or anti-mouse (1:1000 used, Thermo, A31570) secondary antibodies for 1 hour at room temperature. Slides were washed with ice-cold PBS and stained with DAPI diluted in 0.05% Triton-X in PBS for 30 minutes at room temperature. Mounting medium with DAPI (Abcam, ab104139) was used to mount coverslips and left to dry overnight at 4 degrees before imaging.

For neuronal culture, on day 9, attached cells on coverslips were washed once with ice-cold PBS and fixed with 4% paraformaldehyde in PBS for 20 minutes at room temperature. Fixed embryoid bodies were washed once with 0.1% BSA in ice-cold PBS, permeabilized with 0.35% Triton-X for 15 minutes, and blocked for 30 minutes at room temperature. The cells were then incubated with anti-TUBB3 (1ug/mL used, Sigma, T2200-200 ul, lot# 028M4759V) primary antibody at 4 degrees overnight. The cells were washed three times with 0.1% BSA in ice-cold PBS and incubated with anti-rabbit (1:1000 used, Thermo, A31573) secondary antibody for 1 hour at room temperature. The cells were washed with 0.1% BSA in ice-cold PBS and stained with DAPI diluted in 0.05% Triton-X-100 in PBS for 30 minutes at room temperature. Coverslips were mounted to slides using mounting medium with DAPI (Abcam, ab104139). Slides were left to dry overnight at 4 degrees before imaging.

### Neuronal morphology analysis

Neuronal morphology was analysed from confocal microscope images taken with 20X magnification. Images were converted to 8-bit grayscale in ImageJ/Fiji prior to analysis. Neurites were traced semi-automatically using the Fiji plugin, NeuronJ, which allows manual placement of tracing paths with automated centerline detection along neurite processes^57^. The NeuronJ Measure function was used to quantify the total number of neurite tracings and the sum length (μm) of all tracings per day 9 cell.

### RNA-seq

RNA-seq libraries were prepared following a previously described protocol with some modifications^1^. Total RNA was extracted from cells collected on various days of motor neuron differentiation using TRIzol reagent (Thermo, 15596026). NEBNext Poly(A) mRNA Magnetic Isolation Kit (NEB, E7490) was used to remove ribosomal RNA, followed by DNase treatment (Thermo, AM1906). RNA-seq libraries were prepared for sequencing using the NEBNext RNA Library Prep Kit (NEB, E7765) and were sequenced using the Illumina Sequencing Platforms.

### ChIP-seq

Day 6 embryoid bodies containing approximately 30 million cells were collected from culture media, washed with ice-cold PBS, and crosslinked with 1% formaldehyde in PBS for 15 minutes at room temperature^58^. For one replicate of BRG1 ChIP-seq, an additional crosslinking step was performed with 2mM disuccinimidyl glutarate (DSG) for 15 minutes. Cells were lysed, and chromatin pellets were isolated, solubilized, and fragmented into 200-500 bp fragments by sonication^59^. Immunoprecipitation of fragmented chromatin was performed with magnetic beads coupled with Dynabeads Protein A (Invitrogen, 10001D) or Dynabeads Protein G (Invitrogen, 10003D) and antibodies against the protein of interest in 1 mL of 0.5% bovine serum albumin in PBS. Antibodies used in this study are anti-BRG1 (3 ug used, Abcam, ab110641, lot# 1141327-3)) and anti-ARID1B (4 ug used, Abcam, ab57461, lot# 1144777-1), anti-p300 (3 ug used, Abcam, ab275378, lot# 1074860-31 and 3 ug used, Sigma, 05-257, lot# 3213898), anti-CTCF (3 ug used, Sigma, 07-729, lot# 3515588), anti-H3K27ac (3 ug used, Abcam, ab4729, lot# 1059037-6), anti-H3K4me3 (3 ug used, Abcam, ab8580, lot# GR3362386-3), anti-H3K4me1 (3 ug used, Abcam, ab8895, lot# GR3442591-2), and anti-H3K27me3 (3 ug used, Abcam, ab6002, lot# 1035698-1). After washing the beads three times with ChIP wash buffer (500 mM sodium chloride, 50 mM HEPES, 1 mM EDTA, 1% Triton-X, 0.1% sodium deoxycholate, 0.1% sodium dodecyl sulfate), samples were eluted from the magnetic beads in ChIP elution buffer (1% sodium dodecyl sulfate, 10 mM EDTA), reverse crosslinked, and purified. ChIP-seq libraries were prepared for sequencing using the NEBNext DNA Library Prep Kit (NEB, E7645) and sequenced using the Illumina Sequencing Platforms.

### ATAC-seq

ATAC-seq libraries were prepared using the ATAC-seq kit (Active Motif, 53150) and the corresponding protocol. Briefly, day 6 embryoid bodies of differentiated motor neurons were dissociated with trypsin (Thermo, 25300054) into single cells. 75, 000 cells were lysed, and intact nuclei were isolated for tagmentation. Transposed DNA was then column-purified, PCR-amplified, and gel-purified (Qiagen, 28704) before sequencing using the Illumina Sequencing Platforms.

### RNA-seq processing

Raw data files (fastq.gz) were uploaded to Galaxy (https://usegalaxy.org). The raw reads were aligned to the mouse reference genome (mm10) using the Galaxy aligner tool, RNA STAR (version 2.7.11)^60^. The gene annotation file (release M25, GRCm38.p6, comprehensive gene annotation file) was downloaded from GENCODE (www.gencodegenes.org). Then, fragments per kilobase of transcript per million reads (FPKM) were generated with the Galaxy tools Cufflinks package (version 2.2.1.4)^61^. FPKM values were median-normalized across all samples. Prior to converting to logarithmic scales, 0.1 was added to all FPKM values to avoid producing an error for genes with no measured gene expression (FPKM=0). Two biological replicate RNA-seq datasets derived from independent cell differentiations were generated for WT and *Arid1b* KO cells collected on day 0, day 4, and day 6 of differentiation. In addition, we generated one day 5 WT RNA-seq dataset, and the second biological replicate was downloaded from the ENCODE project website (www.encodeproject.org) with GEO accession number GSM2098395^1^.

### Differential gene expression analysis

We performed differential gene expression analysis of WT and *Arid1b* KO cells collected on day 0, day 4, and day 6 separately. For each analysis, we used two biological replicates of each condition derived from independent cell differentiations. First, RNA STAR files were converted to count files using the Galaxy tool featureCounts^62^ and used as input for DESeq2 (version 2.11.40)^63^. FeatureCounts files were also used to generate a normalization file with Galaxy tools Remove Unwanted Variation (version 1.26.0)^64^, which was also used as input for DESeq2. Of the 55,401 genes outputted by DESeq2, only mRNA genes (19,290 genes) were selected for downstream analysis. Differential gene expression analysis of day 0 and day 4 RNA-seq datasets did not detect significant differences between WT and *Arid1b* KO cells. Differential gene expression analysis of day 6 RNA-seq datasets detected significant differences in 12.62% of all annotated protein-coding mRNA genes (2434 of 19,290). The 19,290 mRNA genes were then separated into five groups for analysis including 915 highly downregulated (FDR<0.005, log2 FC < −1.00), 843 highly upregulated (FDR<0.005, log2 FC > 2.00), 676 weakly upregulated (FDR<0.005, 1.00 < log2 FC < 2.00), 16, 425 non-differentially expressed, and 430 unexpressed genes (log2 FC = NC, RC counts = 0) in day 6 *Arid1b* KO cells, compared to WT.

### ChIP-seq and ATAC-seq processing

For ChIP-seq and ATAC-seq datasets, raw data files (fastq.gz) were uploaded to Galaxy America (https://usegalaxy.org). The raw reads were mapped to the mouse reference genome (mm10) using the Galaxy Aligned tool, Bowtie 2 (version 2.5.3)^65^. Bowtie 2 files (.bam) were converted to BigWig file format (bigwig) to visualize peaks using the Integrative Genomics Viewer (IGV; version 2.16.0)^66^. Two biological replicate ChIP-seq datasets from independent differentiations were generated in day 6 WT cells for ARID1B, BRG1, H3K27ac, H3K4me3, H3K4me1, H3K27me3, and p300, and in day 6 *Arid1b* KO cells for ARID1B and H3K27ac. Additionally, we generated one replicate of CTCF ChIP-seq in day 6 WT cells. Publicly available ChIP-seq datasets were obtained from the ENCODE project website (www.encodeproject.org), including day 0 WT ES cells for H3K27ac (GSM1000126), H3K4me3 (GSM1003756), and H3K27me3 (GSM6347289)^67^; day 6 WT motor neurons for CBP (GSM8831504, GSM8831505); and NRF1 (GSM5581267)^47^. Finally, two biological replicate ATAC-seq datasets from independent differentiations were generated in day 6 WT motor neurons.

### ChIP-seq and ATAC-seq analysis

Bowtie files (.bam) were used as input for MACS2 callpeak (version 2.2.9.1)^68^, which was used to identify enriched genomic regions. Two day 6 WT ARID1B ChIP-seq replicates were pooled using MACS2 callpeak, resulting in 22,091 identified peaks (FDR < 0.75). Two day 6 *Arid1b* KO ARID1B ChIP-seq replicates were pooled using MACS2 callpeak, resulting in 4407 identified peaks (FDR < 0.75). Two day 6 WT BRG1 ChIP-seq replicates were pooled using MACS2 callpeak, resulting in 71,744 identified peaks (FDR < 0.05). Within each dataset, we removed ChIP-seq peaks with midpoints that exist <2 kilobase pairs (kb) from one another, keeping the peak with a greater integer score, leaving 18,050 day 6 WT ARID1B ChIP-seq peaks, 3906 day 6 *Arid1b* KO ARID1B ChIP-seq peaks, and 64,667 BRG1 ChIP-seq peaks. Finally, we removed any day 6 WT ARID1B ChIP-seq peaks with a midpoint <2 kb of a day 6 *Arid1b* KO ARID1B ChIP-seq peak (non-specific peaks), which resulted in 16,120 peaks used for downstream analyses.

We defined proximal *cis*-regulatory regions of ChIP-seq peaks as any day 6 WT BRG1 ChIP-seq midpoint <3 kb from an mRNA gene TSS (12,940 BRG1-bound proximal regions). Of these 12,940 regions, we defined ARID1B and BRG1 co-bound proximal regions as any BRG1-bound proximal regions with a day 6 WT ARID1B ChIP-seq peak midpoint within 2 kb. This resulted in 6398/12,940 ARID1B and BRG1 co-bound and 6542/12,940 BRG1-only-bound proximal regions used for downstream analyses. For distal *cis-*regulatory regions analysis, differential binding analysis of day 6 ARID1B and BRG1 ChIP-seq datasets was performed. MACS2 output files (.bed) and Bowtie files (.bam) for two day 6 WT ARID1B ChIP-seq replicates and two day 6 WT BRG1 ChIP-seq replicates were used as input for DiffBind (version 3.12.0)^69^. We defined distal ChIP-seq peaks as any day 6 WT BRG1 ChIP-seq midpoint >3 kb from an mRNA gene TSS (21,613 BRG1-bound distal *cis*-regulatory regions). Of these 21, 613 regions, we defined ARID1B and BRG1 co-bound distal regions as any BRG1-bound distal *cis*-regulatory region with a day 6 WT ARID1B ChIP-seq peak midpoint within 2 kb. This resulted in 4652/21,613 ARID1B and BRG1 co-bound distal regions and 16,961/21,613 BRG1-only-bound distal regions used for downstream analyses.

### Gene ontology (GO) analysis

GO term enrichment analyses were performed using the Gene Ontology Enrichment Analysis and Visualization (GOrilla) system^70^. For GO analyses of gene promoters, the promoters of interest were input as the “target set,” and all 19,290 mRNA genes were input as the “background set”. To remove redundant GO terms, we used the Reduce and Visualize Gene Ontology (REViGO, version 1.8.1)^71^ software with a 0.7 SimRel similarity score based on Resnik’s and Lin’s similarity measures. The top 7 non-redundant GO terms were reported in the figures.

### Motif enrichment analysis

We used the Multiple Em for Motif Elicitation (MEME-ChIP, version 5.5.7)^72^ algorithm for de novo motif discovery. The midpoints of the day 6 WT ATAC-seq peak closest to the day 6 WT BRG1 ChIP-seq peak were used to search for enriched motifs within ARID1B/BRG1 co-bound promoter and distal *cis*-regulatory regions. For MEME-ChIP, 6–12 bp motif widths were searched 100 bp from the midpoints of the identified ATAC-seq peaks. The top enriched *de novo* DNA motifs were searched against a transcription factor binding DNA motif database (JASPAR CORE 2022, non-redundant vertebrate motifs in vivo and in silico) using the motif comparison tool Tomtom (version 5.5.7)^73^, which ranks the DNA motifs in the database and generates an alignment for each significantly matched DNA motif in the database. For the target motif occurrence analysis, we used the Find Individual Motif Occurrences (FIMO, version 5.5.7)^44^. 100 bp from the ATAC-seq midpoints were searched for the occurrence of the target motifs with a matched *p*-value <1.0e-05.

### Box plot analysis

Box plots used to visualize gene expression were generated using median-normalized FPKM values that were log_2_-transformed. Box plots used to visualize ChIP-seq enrichment were generated using normalized reads (i.e., number of reads/genomic location). The ends of the box represent the lower (1^st^) and upper (3^rd^) quartiles, so the box spans the interquartile range. The median is marked by a line inside the box. The whiskers are the two lines outside the box that extend to 1.5 times the interquartile range as the highest and lowest observations. The box plots were generated using KaleidaGraph 5.01 Software (www.synergy.com).

### Statistics and Reproducibility

Statistical analyses were performed using Excel. Comparisons between two groups were conducted using a two-tailed paired Student’s t-test under a null hypothesis because we expected no difference between datasets. Comparisons among three or more groups were analysed by one-way analysis of variance (ANOVA), followed by Tukey’s honestly significant difference (HSD) post hoc test for multiple comparisons when appropriate. All statistical analyses were performed using two biological replicate datasets derived from independent cell differentiations. Data are presented as mean +/− standard deviation. Differences were considered statistically significant at *p* < 0.05.

To test reproducibility of RNA-seq datasets generated in this study, sample-to-sample similarity between RNA-seq biological replicates collected in WT and *Arid1b* KO cells at each stage of differentiation was assessed. Pearson correlation coefficients (*r*) were calculated from the log_2_-transformed FPKM values for 19,290 mRNA genes from two biological replicates derived from independent cell differentiations. The resulting correlation matrices were used to examine the reproducibility of biological replicates (Supplementary Fig. 3). For ChIP-seq and ATAC-seq datasets generated in this study, sample-to-sample similarity between ChIP-seq and ATAC-seq biological replicates collected in WT and *Arid1b* KO cells was assessed. Pearson correlation coefficients (*r*) were calculated from the log_2_-transformed read count values of the 21,601 distal *cis*-regulatory regions defined in **Fig. 4B** from two biological replicates derived from independent cell differentiations. The resulting correlation matrices were used to examine the reproducibility of biological replicates.

## Data availability

Raw and processed sequencing datasets have been deposited in the National Center for Biotechnology Information (NCBI) Gene Expression Omnibus (GEO) under the accession numbers GSE315747 for RNA-seq, GSE317206 for ChIP-seq, and GSE315848 for ATAC-seq. All other relevant data supporting the findings of this study are available within the article and its Supplementary Information files or from the corresponding author upon reasonable request.

## Acknowledgments

We thank Yiming Lu, Guoria Sun, and other members of the Rhee Laboratory for their experimental support and valuable discussions. We thank the Transcription, Chromatin and Epigenetics (TranCE) Community in Toronto for valuable comments. We also thank the Genome Quebec Centre of Expertise and Services and the Centre for Applied Genomics at The Hospital for Sick Children for performing the NGS sequencing.

## Author Contributions

A.I.D.P. and H.S.R. conceived the study and designed experiments and analyses. A.I.D.P. performed most experiments and bioinformatic analyses, including motor neuron differentiation, immunostaining, western blotting, RNA-seq, ChIP-seq, ATAC-seq, cell-cycle assays, and bioinformatic analyses. S.Y.Y. and K.N.M. performed motor neuron differentiation and ChIP-seq. S.Y.Y. performed RNA-seq, and R.J. generated the *Arid1b* KO cell line. A.I.D.P. and H.S.R. co-wrote the manuscript. H.S.R. supervised the overall project. All authors reviewed, contributed to, and approved the final version of the manuscript.

## References

1 Rhee, H. S. et al. Expression of Terminal Effector Genes in Mammalian Neurons Is Maintained by a Dynamic Relay of Transient Enhancers. Neuron 92, 1252–1265 (2016). 10.1016/j.neuron.2016.11.037

2 Hobert, O. Regulatory logic of neuronal diversity: terminal selector genes and selector motifs. Proc Natl Acad Sci U S A 105, 20067–20071 (2008). 10.1073/pnas.0806070105

3 Jaura, R. et al. Extended intergenic DNA contributes to neuron-specific expression of neighboring genes in the mammalian nervous system. Nat Commun 13, 2733 (2022). 10.1038/s41467-022-30192-z

4 Clapier, C. R., Iwasa, J., Cairns, B. R. & Peterson, C. L. Mechanisms of action and regulation of ATP-dependent chromatin-remodelling complexes. Nat Rev Mol Cell Biol 18, 407–422 (2017). 10.1038/nrm.2017.26

5 Bannister, A. J. & Kouzarides, T. Regulation of chromatin by histone modifications. Cell Res 21, 381–395 (2011). 10.1038/cr.2011.22

6 Mizzen, C. A. & Allis, C. D. Linking histone acetylation to transcriptional regulation. Cell Mol Life Sci 54, 6–20 (1998). 10.1007/s000180050121

7 Miller, J. L. & Grant, P. A. The role of DNA methylation and histone modifications in transcriptional regulation in humans. Subcell Biochem 61, 289–317 (2013). 10.1007/978-94-007-4525-4_13

8 Alfert, A., Moreno, N. & Kerl, K. The BAF complex in development and disease. Epigenetics Chromatin 12, 19 (2019). 10.1186/s13072-019-0264-y

9 Sokpor, G., Castro-Hernandez, R., Rosenbusch, J., Staiger, J. F. & Tuoc, T. ATP-Dependent Chromatin Remodeling During Cortical Neurogenesis. Front Neurosci 12, 226 (2018). 10.3389/fnins.2018.00226

10 Alver, B. H. et al. The SWI/SNF chromatin remodelling complex is required for maintenance of lineage specific enhancers. Nat Commun 8, 14648 (2017). 10.1038/ncomms14648

11 Lessard, J. et al. An essential switch in subunit composition of a chromatin remodeling complex during neural development. Neuron 55, 201–215 (2007). 10.1016/j.neuron.2007.06.019

12 Olave, I., Wang, W., Xue, Y., Kuo, A. & Crabtree, G. R. Identification of a polymorphic, neuron-specific chromatin remodeling complex. Genes Dev 16, 2509–2517 (2002). 10.1101/gad.992102

13 Kaitlin N. Montanera, A. J. L., Woobin Yun, Alyssa Ialongo Del Prete, Ssu-Yu Yeh, Inkyung Jung, Ho Sung Rhee. CREST functions as a molecular bridge between neuron-specific enhancers and epigenetic regulators in mature motor neurons. Nature Communications In press (2026). 10.1038/s41467-026-78396-x

14 Pagliaroli, L. et al. Inability to switch from ARID1A-BAF to ARID1B-BAF impairs exit from pluripotency and commitment towards neural crest formation in ARID1B-related neurodevelopmental disorders. Nat Commun 12, 6469 (2021). 10.1038/s41467-021-26810-x

15 Flores-Alcantar, A., Gonzalez-Sandoval, A., Escalante-Alcalde, D. & Lomelí, H. Dynamics of expression of ARID1A and ARID1B subunits in mouse embryos and in cells during the cell cycle. Cell Tissue Res 345, 137–148 (2011). 10.1007/s00441-011-1182-x

16 Hoyer, J. et al. Haploinsufficiency of ARID1B, a member of the SWI/SNF-a chromatin-remodeling complex, is a frequent cause of intellectual disability. Am J Hum Genet 90, 565–572 (2012). 10.1016/j.ajhg.2012.02.007

17 Ellegood, J. et al. Neuroanatomy and behavior in mice with a haploinsufficiency of AT-rich interactive domain 1B (ARID1B) throughout development. Mol Autism 12, 25 (2021). 10.1186/s13229-021-00432-y

18 van der Sluijs, P. J. et al. -related disorder in 87 adults: Natural history and self-sustainability. Genet Med Open 2, 101873 (2024). 10.1016/j.gimo.2024.101873

19 Jung, E. M. et al. Arid1b haploinsufficiency disrupts cortical interneuron development and mouse behavior. Nat Neurosci 20, 1694–1707 (2017). 10.1038/s41593-017-0013-0

20 Ka, M., Chopra, D. A., Dravid, S. M. & Kim, W. Y. Essential Roles for ARID1B in Dendritic Arborization and Spine Morphology of Developing Pyramidal Neurons. J Neurosci 36, 2723–2742 (2016). 10.1523/JNEUROSCI.2321-15.2016

21 Martins-Costa, C. et al. ARID1B controls transcriptional programs of axon projection in an organoid model of the human corpus callosum. Cell Stem Cell 31, 866–885.e814 (2024). 10.1016/j.stem.2024.04.014

22 Stifani, N. Motor neurons and the generation of spinal motor neuron diversity. Front Cell Neurosci 8, 293 (2014). 10.3389/fncel.2014.00293

23 Jessell, T. M. Neuronal specification in the spinal cord: inductive signals and transcriptional codes. Nat Rev Genet 1, 20–29 (2000). 10.1038/35049541

24 Scardigli, R., Schuurmans, C., Gradwohl, G. & Guillemot, F. Crossregulation between Neurogenin2 and pathways specifying neuronal identity in the spinal cord. Neuron 31, 203–217 (2001). 10.1016/s0896-6273(01)00358-0

25 Sim, J. C., White, S. M. & Lockhart, P. J. ARID1B-mediated disorders: Mutations and possible mechanisms. Intractable Rare Dis Res 4, 17–23 (2015). 10.5582/irdr.2014.01021

26 Lee, S. et al. Fusion protein Isl1-Lhx3 specifies motor neuron fate by inducing motor neuron genes and concomitantly suppressing the interneuron programs. Proc Natl Acad Sci U S A 109, 3383–3388 (2012). 10.1073/pnas.1114515109

27 Arber, S. et al. Requirement for the homeobox gene Hb9 in the consolidation of motor neuron identity. Neuron 23, 659–674 (1999). 10.1016/s0896-6273(01)80026-x

28 Gleeson, J. G., Lin, P. T., Flanagan, L. A. & Walsh, C. A. Doublecortin is a microtubule-associated protein and is expressed widely by migrating neurons. Neuron 23, 257–271 (1999). 10.1016/s0896-6273(00)80778-3

29 Blockus, H. & Chédotal, A. Slit-Robo signaling. Development 143, 3037–3044 (2016). 10.1242/dev.132829

30 Hill, G. W. et al. Netrin-1-mediated axon guidance in mouse embryonic stem cells overexpressing neurogenin-1. Stem Cells Dev 21, 2827–2837 (2012). 10.1089/scd.2011.0437

31 Deloulme, J. C. et al. Microtubule-associated protein 6 mediates neuronal connectivity through Semaphorin 3E-dependent signalling for axonal growth. Nat Commun 6, 7246 (2015). 10.1038/ncomms8246

32 Taft, C. E. & Turrigiano, G. G. PSD-95 promotes the stabilization of young synaptic contacts. Philos Trans R Soc Lond B Biol Sci 369, 20130134 (2014). 10.1098/rstb.2013.0134

33 De Bertier, S. et al. MAPT mutations in amyotrophic lateral sclerosis: clinical, neuropathological and functional insights. J Neurol 272, 272 (2025). 10.1007/s00415-025-13007-1

34 Christ, A. et al. LRP2 is an auxiliary SHH receptor required to condition the forebrain ventral midline for inductive signals. Dev Cell 22, 268–278 (2012). 10.1016/j.devcel.2011.11.023

35 Dashti, P. et al. G-protein coupled receptor 5C (GPRC5C) is required for osteoblast differentiation and responds to EZH2 inhibition and multiple osteogenic signals. Bone 176, 116866 (2023). 10.1016/j.bone.2023.116866

36 Jackson, R. J., Hyman, B. T. & Serrano-Pozo, A. Multifaceted roles of APOE in Alzheimer disease. Nat Rev Neurol 20, 457–474 (2024). 10.1038/s41582-024-00988-2

37 Pellarin, I. et al. Cyclin-dependent protein kinases and cell cycle regulation in biology and disease. Signal Transduct Target Ther 10, 11 (2025). 10.1038/s41392-024-02080-z

38 Lin, X. et al. The Role of CDCA2 in tumor genesis, prognosis and future treatments. Eur J Cancer 211, 114308 (2024). 10.1016/j.ejca.2024.114308

39 Heintzman, N. D. et al. Distinct and predictive chromatin signatures of transcriptional promoters and enhancers in the human genome. Nat Genet 39, 311–318 (2007). 10.1038/ng1966

40 Glancy, E., Choy, N. & Eckersley-Maslin, M. A. Bivalent chromatin: a developmental balancing act tipped in cancer. Biochem Soc Trans 52, 217–229 (2024). 10.1042/BST20230426

41 Sánchez-Sarasúa, S. et al. IRS1 expression in hippocampus is age-dependent and is required for mature spine maintenance and neuritogenesis. Mol Cell Neurosci 118, 103693 (2022). 10.1016/j.mcn.2021.103693

42 Vukojevic, V. et al. Evolutionary conserved role of neural cell adhesion molecule-1 in memory. Transl Psychiatry 10, 217 (2020). 10.1038/s41398-020-00899-y

43 Wang, X. et al. Two related ARID family proteins are alternative subunits of human SWI/SNF complexes. Biochem J 383, 319–325 (2004). 10.1042/BJ20040524

44 Grant, C. E., Bailey, T. L. & Noble, W. S. FIMO: scanning for occurrences of a given motif. Bioinformatics 27, 1017–1018 (2011). 10.1093/bioinformatics/btr064

45 Issac, J., Raveendran, P. S. & Das, A. V. RFX1: a promising therapeutic arsenal against cancer. Cancer Cell Int 21, 253 (2021). 10.1186/s12935-021-01952-6

46 Schwab, K. et al. Multi-omics analysis identifies RFX7 targets involved in tumor suppression and neuronal processes. Cell Death Discov 9, 80 (2023). 10.1038/s41420-023-01378-1

47 Liu, S. et al. NRF1 association with AUTS2-Polycomb mediates specific gene activation in the brain. Mol Cell 81, 4757 (2021). 10.1016/j.molcel.2021.10.023

48 Beamish, H. et al. Cyclin A/cdk2 regulates adenomatous polyposis coli-dependent mitotic spindle anchoring. J Biol Chem 284, 29015–29023 (2009). 10.1074/jbc.M109.042820

49 Geisert, E. E. & Frankfurter, A. The neuronal response to injury as visualized by immunostaining of class III beta-tubulin in the rat. Neurosci Lett 102, 137–141 (1989). 10.1016/0304-3940(89)90068-2

50 Moffat, J. J. et al. Differential roles of ARID1B in excitatory and inhibitory neural progenitors in the developing cortex. Sci Rep 11, 3856 (2021). 10.1038/s41598-021-82974-y

51 Wu, J. I. Diverse functions of ATP-dependent chromatin remodeling complexes in development and cancer. Acta Biochim Biophys Sin (Shanghai) 44, 54–69 (2012). 10.1093/abbs/gmr099

52 Patsialou, A., Wilsker, D. & Moran, E. DNA-binding properties of ARID family proteins. Nucleic Acids Res 33, 66–80 (2005). 10.1093/nar/gki145

53 Sherwood, R. I. et al. Discovery of directional and nondirectional pioneer transcription factors by modeling DNase profile magnitude and shape. Nat Biotechnol 32, 171–178 (2014). 10.1038/nbt.2798

54 Hodges, C., Kirkland, J. G. & Crabtree, G. R. The Many Roles of BAF (mSWI/SNF) and PBAF Complexes in Cancer. Cold Spring Harb Perspect Med 6 (2016). 10.1101/cshperspect.a026930

55 Euskirchen, G. M. et al. Diverse roles and interactions of the SWI/SNF chromatin remodeling complex revealed using global approaches. PLoS Genet 7, e1002008 (2011). 10.1371/journal.pgen.1002008

56 Ran, F. A. et al. Genome engineering using the CRISPR-Cas9 system. Nat Protoc 8, 2281–2308 (2013). 10.1038/nprot.2013.143

57 Meijering, E. et al. Design and validation of a tool for neurite tracing and analysis in fluorescence microscopy images. Cytometry A 58, 167–176 (2004). 10.1002/cyto.a.20022

58 Yeh, S. Y. & Rhee, H. S. The ChIP-Exo Method to Identify Genomic Locations of DNA-Binding Proteins at Near Single Base-Pair Resolution. Methods Mol Biol 2599, 33–48 (2023). 10.1007/978-1-0716-2847-8_4

59 Montanera, K. N. & Rhee, H. S. High-Resolution Mapping of Protein-DNA Interactions in Mouse Stem Cell-Derived Neurons using Chromatin Immunoprecipitation-Exonuclease (ChIP-Exo). J Vis Exp (2020). 10.3791/61124

60 Dobin, A. et al. STAR: ultrafast universal RNA-seq aligner. Bioinformatics 29, 15–21 (2013). 10.1093/bioinformatics/bts635

61 Trapnell, C. et al. Transcript assembly and quantification by RNA-Seq reveals unannotated transcripts and isoform switching during cell differentiation. Nat Biotechnol 28, 511–515 (2010). 10.1038/nbt.1621

62 Liao, Y., Smyth, G. K. & Shi, W. featureCounts: an efficient general purpose program for assigning sequence reads to genomic features. Bioinformatics 30, 923–930 (2014). 10.1093/bioinformatics/btt656

63 Love, M. I., Huber, W. & Anders, S. Moderated estimation of fold change and dispersion for RNA-seq data with DESeq2. Genome Biol 15, 550 (2014). 10.1186/s13059-014-0550-8

64 Risso, D., Ngai, J., Speed, T. P. & Dudoit, S. Normalization of RNA-seq data using factor analysis of control genes or samples. Nat Biotechnol 32, 896–902 (2014). 10.1038/nbt.2931

65 Langmead, B. & Salzberg, S. L. Fast gapped-read alignment with Bowtie 2. Nat Methods 9, 357–359 (2012). 10.1038/nmeth.1923

66 Thorvaldsdóttir, H., Robinson, J. T. & Mesirov, J. P. Integrative Genomics Viewer (IGV): high-performance genomics data visualization and exploration. Brief Bioinform 14, 178–192 (2013). 10.1093/bib/bbs017

67 Consortium, E. P. An integrated encyclopedia of DNA elements in the human genome. Nature 489, 57–74 (2012). 10.1038/nature11247

68 Zhang, Y. et al. Model-based analysis of ChIP-Seq (MACS). Genome Biol 9, R137 (2008). 10.1186/gb-2008-9-9-r137

69 Wu, D. Y., Bittencourt, D., Stallcup, M. R. & Siegmund, K. D. Identifying differential transcription factor binding in ChIP-seq. Front Genet 6, 169 (2015). 10.3389/fgene.2015.00169

70 Eden, E., Navon, R., Steinfeld, I., Lipson, D. & Yakhini, Z. GOrilla: a tool for discovery and visualization of enriched GO terms in ranked gene lists. BMC Bioinformatics 10, 48 (2009). 10.1186/1471-2105-10-48

71 Supek, F., Bosnjak, M., Skunca, N. & Smuc, T. REVIGO summarizes and visualizes long lists of gene ontology terms. PLoS One 6, e21800 (2011). 10.1371/journal.pone.0021800

72 Machanick, P. & Bailey, T. L. MEME-ChIP: motif analysis of large DNA datasets. Bioinformatics 27, 1696–1697 (2011). 10.1093/bioinformatics/btr189

73 Gupta, S., Stamatoyannopoulos, J. A., Bailey, T. L. & Noble, W. S. Quantifying similarity between motifs. Genome Biol 8, R24 (2007). 10.1186/gb-2007-8-2-r24

