## Supplementary Figure for "The BAF chromatin remodeling subunit ARID1B coordinates cell-cycle exit and motor neuron differentiation"

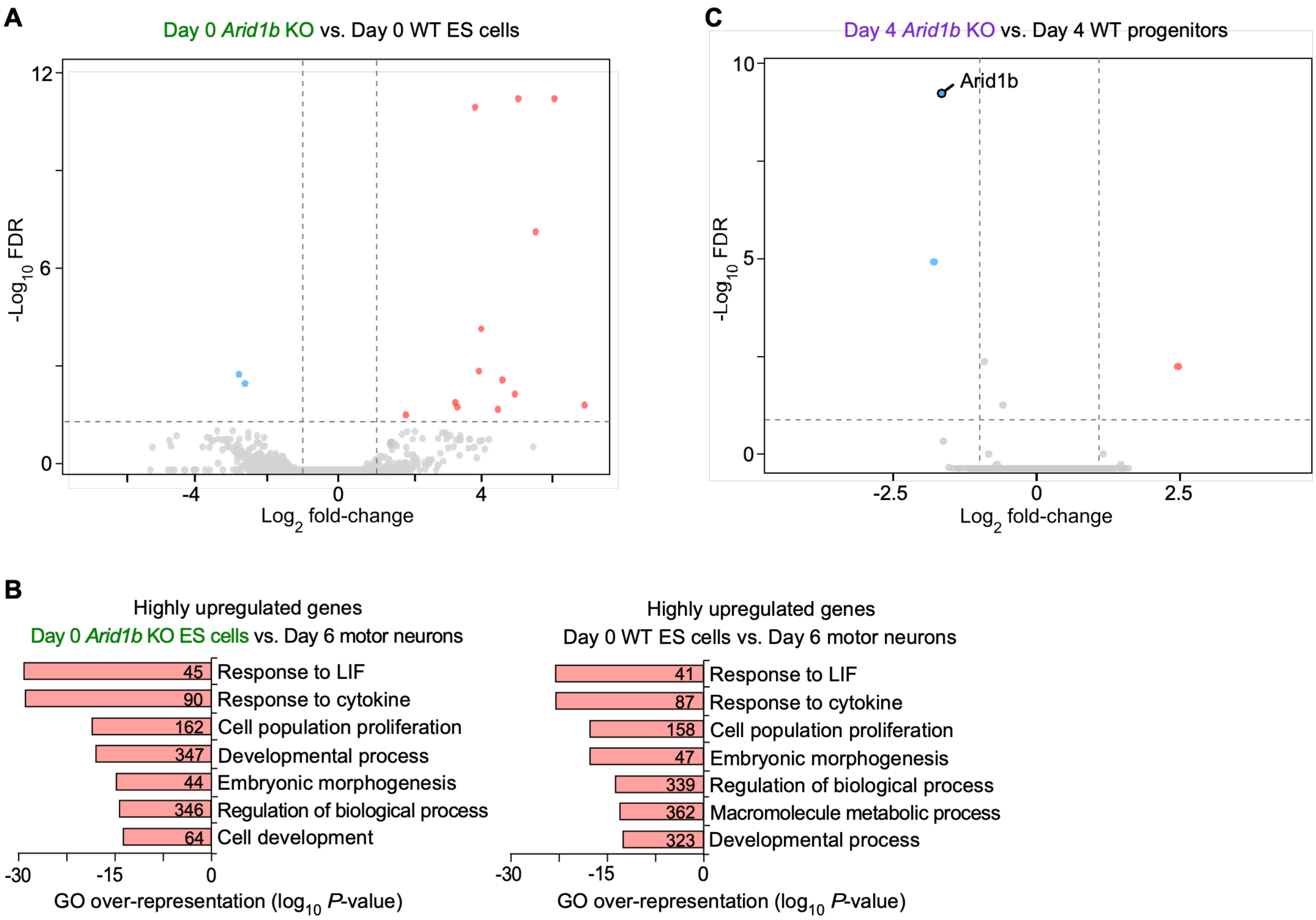


**Supplementary Fig. 1** | ***Arid1b* knockout does not substantially alter global gene expression in day 0 ES cells or day 4 neural progenitors.**

**A.** Volcano plot of RNA-seq data for all 19,290 mRNA genes comparing day 0 *Arid1b* KO and day 0 WT ES cells. Differentially expressed genes were identified using DESeq2 and classified as downregulated (log2 fold change [FC] < −1.00, FDR < 0.005) or upregulated (log2 FC > 1.00, FDR < 0.005). Data from two biological replicates derived from independent cell differentiations were analyzed (Supplementary Data 2).

**B.** Gene Ontology analysis of the top 5% of genes with the highest expression in day 0 *Arid1b* KO (*left*) and day 0 WT ES cells (*right*) compared with day 6 WT motor neurons, measured by RNA-seq. The seven most significantly enriched, nonredundant GO terms are shown. Numbers within bars represent the number of genes associated with each GO term (Supplementary Data 5, 6).

**C.** Volcano plot of RNA-seq data for all 19,290 mRNA genes comparing day 4 *Arid1b* KO and day 4 WT neural progenitors. Differentially expressed genes were identified using DESeq2 and classified as downregulated (log2 fold change [FC] < −1.00, FDR < 0.005) or upregulated (log2 FC > 1.00, FDR < 0.005). Data from two biological replicates derived from independent cell differentiations were analyzed (Supplementary Data 2).

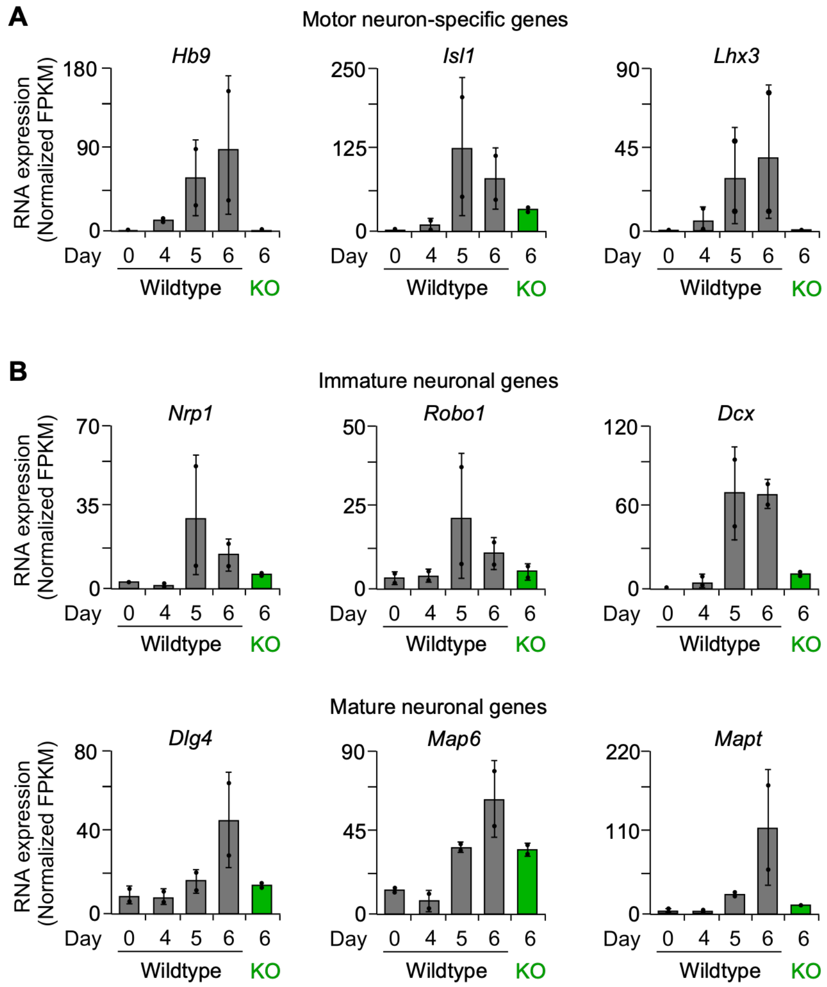


**Supplementary Fig. 2 | *Arid1b* knockout impairs neuronal gene expression during motor neuron differentiation.**

**A.** mRNA expression of motor neuron-specific genes (*Hb9, Isl1,* and *Lhx3*) measured by RNA-seq in WT cells on days 0, 4, 5, and 6 and in day 6 *Arid1b* KO cells. FPKM values from two biological replicates derived from independent differentiations were median-normalized and averaged (Supplementary Data 3, 4). Data points represent median-normalized FPKM values from individual biological replicates ($n$ = 2); error bars represent the standard deviation.

**B.** mRNA expression of immature neuronal genes (*Nrp1, Robo1,* and *Dcx*) and mature neuronal genes (*Dlg4, Map6,* and *Mapt*), measured and presented as described in panel **A**.

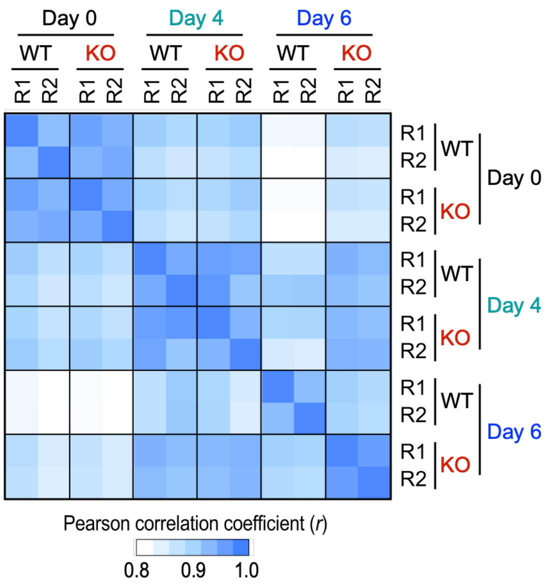


**Supplementary Fig. 3 | Pearson correlation analysis of RNA-seq datasets demonstrates reproducibility across biological replicates.**

Pearson correlation coefficients (*r*) between RNA-seq samples generated in this study were calculated using $log_2$-transformed FPKM values across all annotated mRNA genes (*n* = 19,290) from two biological replicates derived from independent cell differentiations (R1 and R2). The heatmap shows pairwise correlations among all examined samples (Supplementary Data 8).

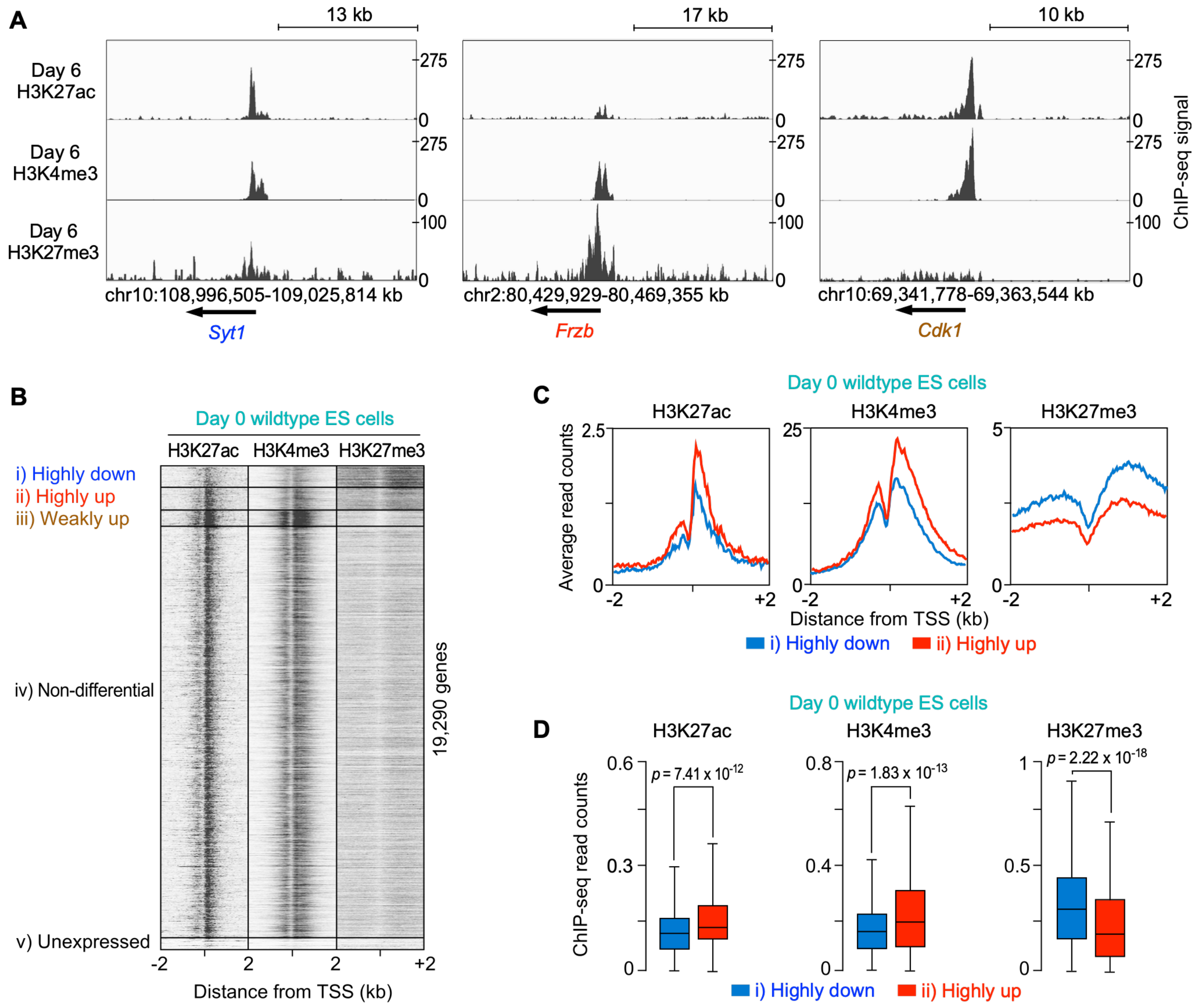


**Supplementary Fig. 4** | **Differentially expressed gene promoters exhibit inverse histone modification patterns in day 0 ES cells relative to day 6 motor neurons.**

**A.** Representative genomic loci for the highly downregulated *Syt1* (synaptotagmin-1), highly upregulated *Frzb* (frizzled-related protein), and weakly upregulated *Cdk1* (cyclin-dependent kinase 1) genes, identified in **Fig. 2A**. H3K27ac, H3K4me3, and H3K27me3 enrichment was measured by ChIP-seq in day 6 WT motor neurons.

**B.** Heatmaps of H3K27ac, H3K4me3, and H3K27me3 ChIP-seq signals in day 0 WT ES cells (Supplementary Data 2). Signals were plotted relative to the transcription start sites (TSSs) of all 19,290 mRNA genes, ordered and grouped as in **Fig. 2A**.

**C.** Composite plots of H3K27ac, H3K4me3, and H3K27me3 ChIP-seq signals at promoters of highly downregulated (Group *i*, blue) and highly upregulated (Group *ii*, red) genes in day 0 WT ES cells, as defined in panel **B**.

**D.** Box plots comparing H3K27ac, H3K4me3, and H3K27me3 ChIP-seq signals at promoters of highly downregulated (Group *i*) and highly upregulated (Group *ii*) genes in day 0 WT ES cells (Supplementary Data 2). Boxes indicate the interquartile range (IQR), with center lines indicating the median and whiskers extending to 1.5 × IQR (Supplementary Data 4). *P*-values were calculated using two-sided paired *t*-tests ($n = 1$).

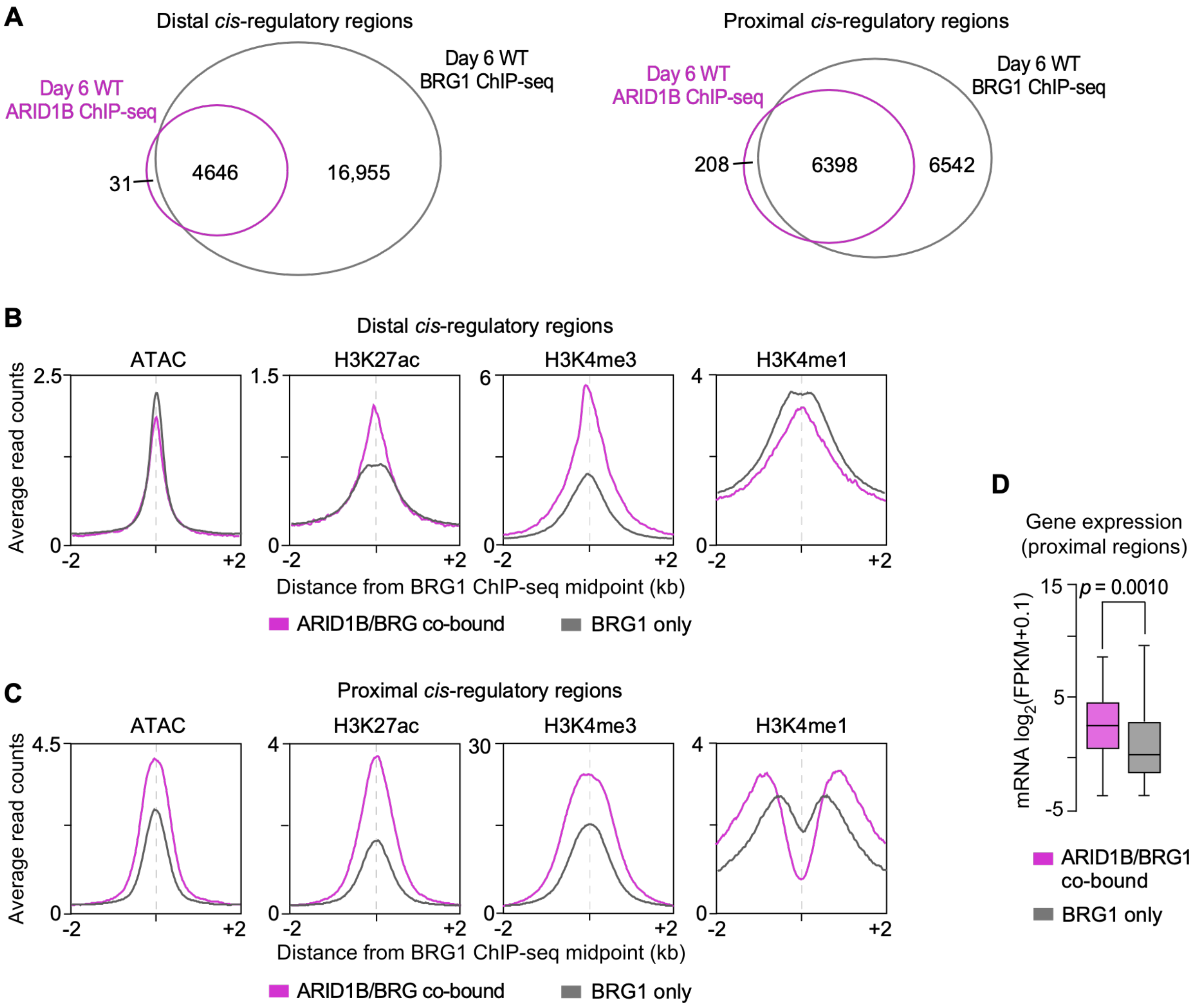


**Supplementary Fig. 5 | ARID1B/BRG1 co-bound *cis*-regulatory regions exhibit distinct epigenetic signatures compared with BRG1-only bound regions.**

**A.** *Left:* Venn diagram comparing ARID1B and BRG1 ChIP-seq peaks at distal *cis*-regulatory regions in day 6 WT motor neurons. *Right*: Venn diagram comparing ARID1B and BRG1 ChIP-seq peaks at proximal *cis*-regulatory regions. A total of 96.85% of ARID1B-bound promoter peaks (6,398 of 6,606 regions) were located within 2 kb of BRG1 ChIP-seq peaks.

**B.** Composite plots of ATAC-seq signals and H3K27ac, H3K4me3, and H3K4me1 ChIP-seq signals at ARID1B/BRG1 co-bound and BRG1-only bound distal *cis*-regulatory regions corresponding to the heatmaps shown in **Fig. 4C**.

**C.** Composite plots of ATAC-seq signals and H3K27ac, H3K4me3, and H3K4me1 ChIP-seq signals at ARID1B/BRG1 co-bound and BRG1-only bound proximal *cis*-regulatory regions, displayed as in panel **B**.

**D.** Box plots showing RNA expression ($\log_2$[FPKM + 0.1]) of genes associated with ARID1B/BRG1 co-bound and BRG1-only bound proximal *cis*-regulatory regions shown in panel **C** (Supplementary Data 4). RNA expression data from two biological replicates from independent differentiations were median-normalized, averaged, and $\log_2$-transformed ($n$ = 2). *P*-values were calculated using a two-sided paired *t*-test.

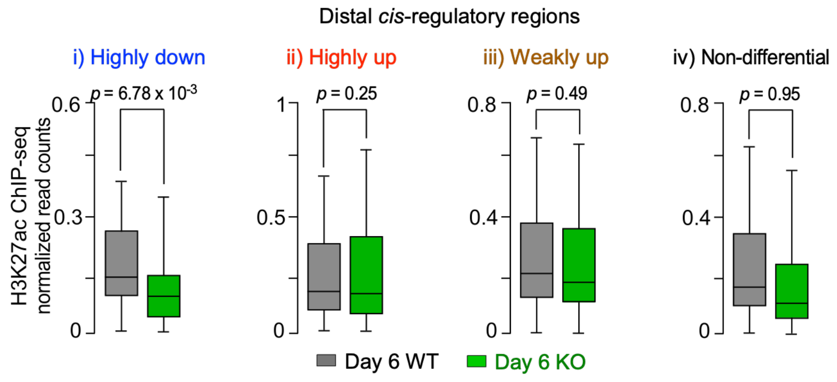


**Supplementary Fig. 6 | H3K27ac enrichment at ARID1B/BRG1 co-bound distal *cis*-regulatory regions.**

Box plots comparing H3K27ac, H3K4me3, and H3K27me3 ChIP-seq signals at ARID1B/BRG1 co-bound distal *cis*-regulatory regions associated with the gene groups shown in **Fig. 4D** (Supplementary Data 4). Boxes indicate the interquartile range (IQR), with center lines indicating the median and whiskers extending to 1.5 × IQR. *P*-values were calculated using two-sided paired *t*-tests ($n = 2$).

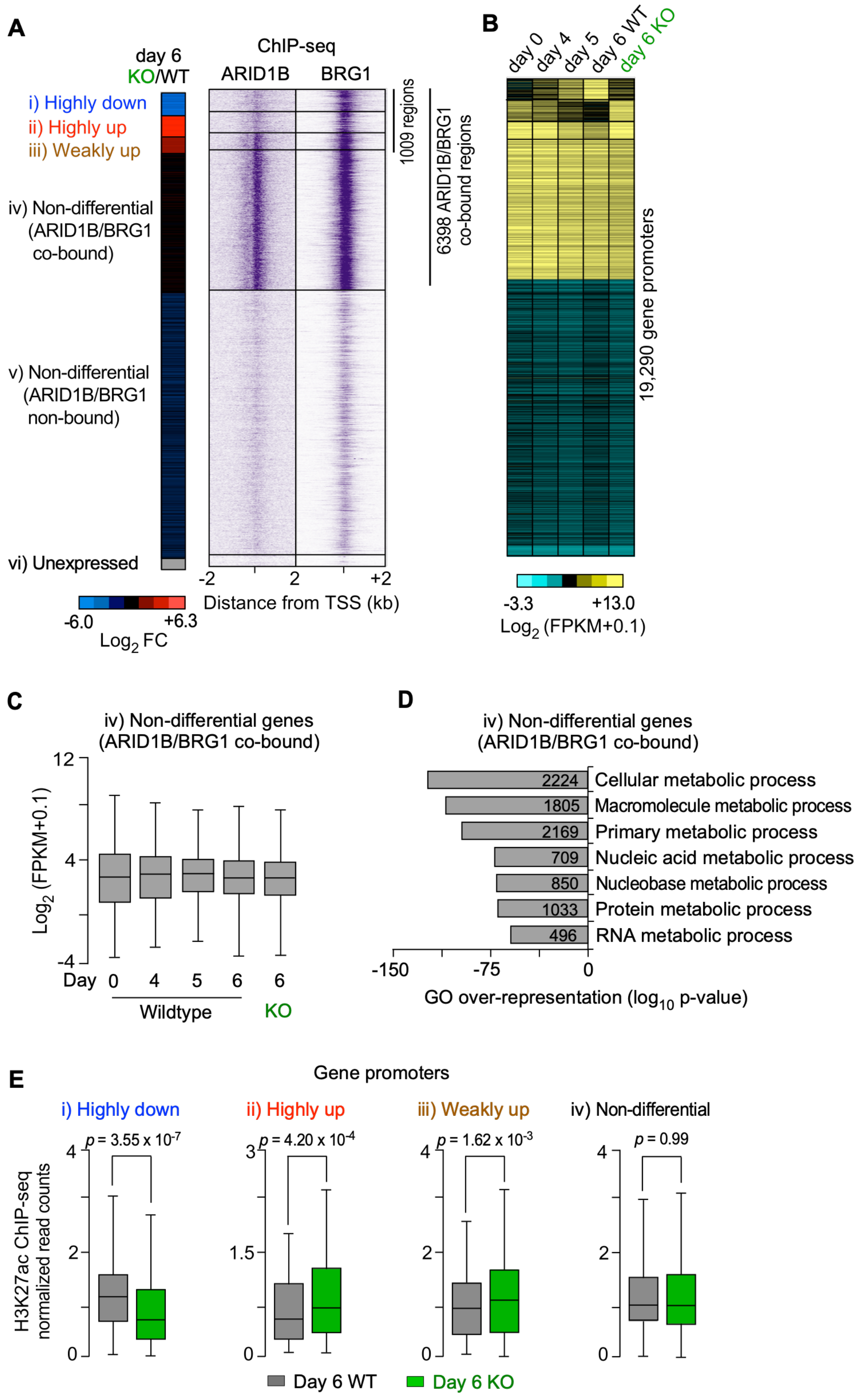


**Supplementary Fig. 7 | ARID1B/BRG1 co-bind promoters of differentially and non-differentially expressed genes.**

**A.** *Left:* Heatmap of differential gene expression ($\log_2$ fold change [FC]) for all 19,290 mRNA genes in day 6 *Arid1b* KO cells compared with day 6 WT motor neurons, measured by RNA-seq, as in **Fig. 2A**, with non-differentially expressed genes further classified according to ARID1B/BRG1 promoter co-occupancy (Groups *iv* and *v*). *Right:* Heatmaps of ARID1B and

BRG1 ChIP-seq signals in day 6 WT motor neurons. All signals were plotted relative to the transcription start sites (TSSs) of all 19,290 mRNA genes and grouped and ordered as displayed on the *left*.

**B.** Heatmap of RNA expression ($log_2$[FPKM + 0.1]) of all 19,290 mRNA genes in WT cells on days 0, 4, 5, and 6 and in day 6 Arid1b KO cells, measured by RNA-seq. All 19,290 mRNA genes were grouped and ordered as in panel **A**.

**C.** Box plots showing RNA expression ($log_2$[FPKM + 0.1]) across differentiation stages for non-differentially expressed genes whose promoters are co-bound by ARID1B and BRG1 (Group *iv*), as shown in panel **B** (Supplementary Data 4). RNA expression data from two biological replicates derived from independent cell differentiations were median-normalized, averaged, and $log_2$-transformed ($n$ = 2). Boxes indicate the interquartile range (IQR), with center lines indicating the median and whiskers extending to 1.5 × IQR.

**D.** Gene Ontology analysis of non-differentially expressed genes whose promoters are co-bound by ARID1B and BRG1 (Group iv), as shown in panel **B**. The seven most significantly enriched, nonredundant GO terms are shown. Numbers within bars represent the number of genes associated with each GO term (Supplementary Data 5).

**E.** Box plots comparing H3K27ac, H3K4me3, and H3K27me3 ChIP-seq signals at ARID1B/BRG1 co-bound promoters associated with the gene groups shown in **Fig. 4E** (Supplementary Data 4). Boxes indicate the interquartile range (IQR), with center lines indicating the median and whiskers extending to 1.5 × IQR. *P*-values were calculated using two-sided paired *t*-tests ($n$ = 2).

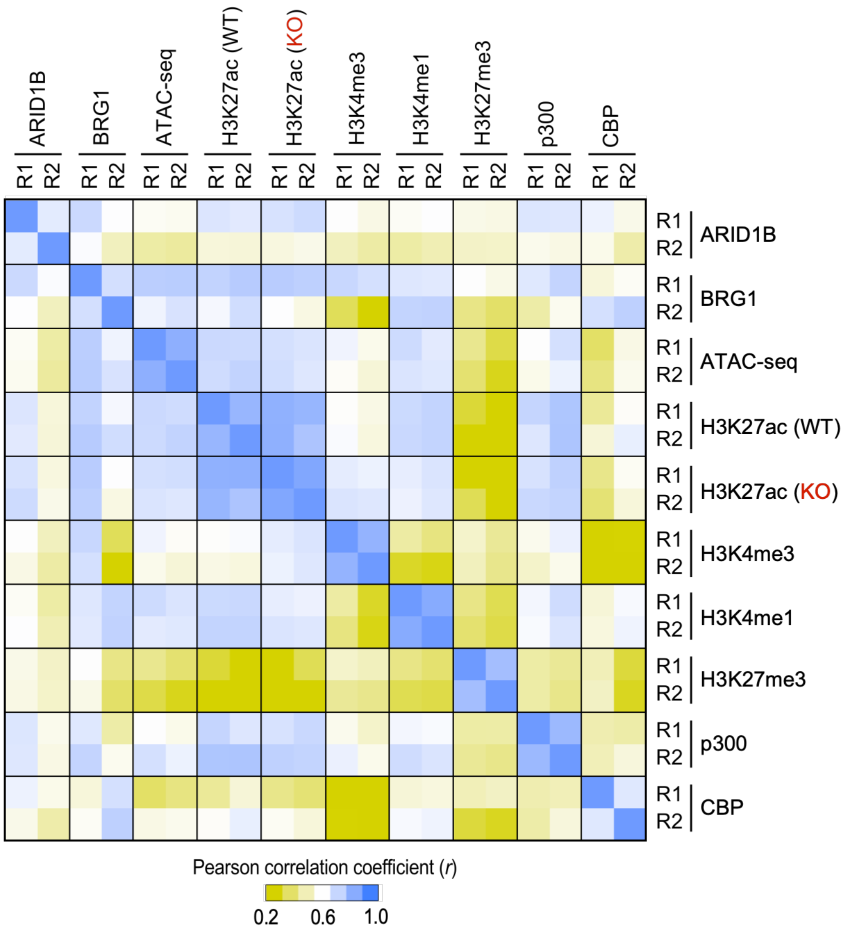


**Supplementary Fig. 8** | **Pearson correlation analysis of ChIP-seq and ATAC-seq datasets demonstrates reproducibility across biological replicates.**

Pearson correlation coefficients (*r*) between ChIP-seq and ATAC-seq samples generated in this study were calculated using $log_2$-transformed read counts across 21,601 distal *cis*-regulatory regions defined in **Fig. 4B** from two biological replicates derived from independent cell differentiations (R1 and R2). The heatmap shows pairwise correlations among all examined chromatin-associated signals (Supplementary Data 8).

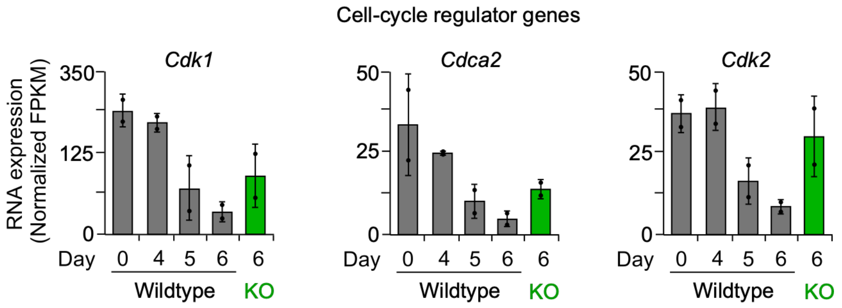


**Supplementary Fig. 9** | **Repression of mitotic cell-cycle genes during motor neuron differentiation requires ARID1B.**

mRNA expression of mitotic cell-cycle genes (*Cdk1, Cdca2,* and *Cdk2*) was measured by RNA-seq in WT cells on days 0, 4, 5, and 6 and in day 6 *Arid1b* KO cells. FPKM values from two biological replicates derived from independent differentiations were median-normalized and averaged (Supplementary Data 3, 4). Data points represent median-normalized FPKM values from individual biological replicates ($n$ = 2); error bars represent the standard deviation.
